# Integrative Mapping of Pre-existing Immune Landscapes for Vaccine Response Prediction

**DOI:** 10.1101/2025.01.22.634302

**Authors:** Stephanie Hao, Ivan Tomic, Benjamin B Lindsey, Ya Jankey Jagne, Katja Hoschler, Adam Meijer, Juan Manuel Carreño Quiroz, Philip Meade, Kaori Sano, Chikondi Peno, André G Costa-Martins, Debby Bogaert, Beate Kampmann, Helder Nakaya, Florian Krammer, Thushan I de Silva, Adriana Tomic

## Abstract

Predicting individual vaccine responses remains a significant challenge due to the complexity and variability of immune processes. To address this gap, we developed *immunaut*, an open-source, data-driven framework implemented as an R package specifically designed for all systems vaccinologists seeking to analyze and predict immunological outcomes across diverse vaccination settings. Leveraging one of the most comprehensive live attenuated influenza vaccine (LAIV) datasets to date - 244 Gambian children enrolled in a phase 4 immunogenicity study - *immunaut* integrates humoral, mucosal, cellular, transcriptomic, and microbiological parameters collected before and after vaccination, providing an unprecedentedly holistic view of LAIV-induced immunity. Through advanced dimensionality reduction, clustering, and predictive modeling, immunaut identifies distinct immunophenotypic responder profiles and their underlying baseline determinants. In this study, *immunaut* delineated three immunophenotypes: (1) CD8 T-cell responders, marked by strong baseline mucosal immunity and extensive prior influenza virus exposure that boosts memory CD8 T-cell responses, without generating influenza virus-specific antibody responses; (2) Mucosal responders, characterized by pre-existing systemic influenza A virus immunity (specifically to H3N2) and stable epithelial integrity, leading to potent mucosal IgA expansions and subsequent seroconversion to influenza B virus; and (3) Systemic, broad influenza A virus responders, who start with relatively naive immunity and leverage greater initial viral replication to drive broad systemic antibody responses against multiple influenza A virus variants beyond those included in the LAIV vaccine. By integrating pathway-level analysis, model-derived contribution scores, and hierarchical decision rules, *immunaut* elucidates how distinct immunological landscapes shape each response trajectory and how key baseline features, including pre-existing immunity, mucosal preparedness, and cellular support, dictate vaccine outcomes. Collectively, these findings emphasize the power of integrative, predictive frameworks to advance precision vaccinology, and highlight *immunaut* as a versatile, community-available resource for optimizing immunization strategies across diverse populations and vaccine platforms.

**One-Sentence Summary:** Using one of the most comprehensive LAIV datasets compiled to date, *immunaut*, an integrative machine learning framework, identifies distinct immunophenotypic responder groups shaped by baseline immune landscapes, advancing precision vaccinology and guiding more effective, personalized immunization strategies.

**Graphical abstract:** *Immunaut*, an automated framework for mapping and predicting vaccine response immunotypes. Step 1 outlines the identification of vaccine response outcomes using pre- and post-vaccination data integration across immune features, including antibodies, flu-specific T-cells, and immunophenotyping at mucosal and systemic sites. Clustering methods define the vaccine response landscape, stability, and validation through t-SNE-based visualization. Step 2 leverages an automated machine learning modeling approach, to enhance the accuracy and interpretability of vaccine response predictions, enabling stratification and targeted intervention strategies for personalized vaccine immunogenicity.

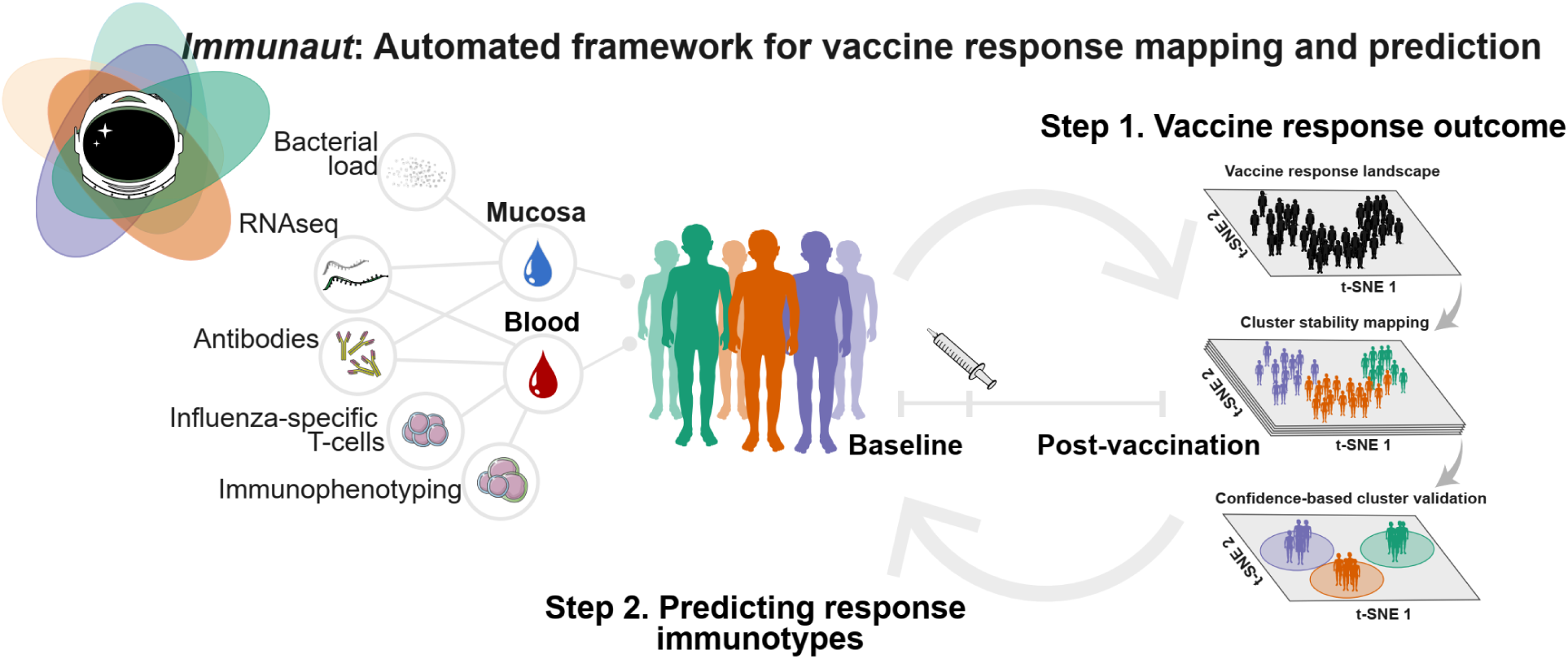

## Introduction

Understanding the ability of a vaccine to elicit an effective immune response, *i.e*., vaccine immunogenicity, is fundamental to guiding vaccination programs and advancing public health initiatives^1^. Traditional evaluation methods often measure humoral and cellular immunity in isolation, overlooking the intricate interplay between different immune compartments^2–4^. Although emerging high-dimensional profiling technologies now enable more holistic assessments^3^, comprehensive, large-scale evaluations that simultaneously capture systemic, mucosal, and cellular immune responses remain rare. This scarcity of broad, systems-level evaluations has posed a significant barrier to fully understanding and predicting vaccine-induced immunity, especially for complex vaccines such as live attenuated influenza vaccine (LAIV), which engages multiple arms of the adaptive immune system concurrently^5, 6^.

Here, we bridge this gap by leveraging a uniquely extensive LAIV dataset - one that, to our knowledge, surpasses previous studies in both depth and breadth. Drawing on a cohort of 244 children aged 24-59 months in The Gambia enrolled in a phase 4 immunogenicity study^7–9^, we integrated humoral, cellular, and mucosal responses with detailed baseline clinical and immunological measurements. Unlike prior work focusing on narrower immune readouts, our dataset encompasses multiple serum antibody and T-cell responses, mucosal IgA, transcriptomic profiles, and microbiological assessments collected before and after vaccination. This unprecedented scope provides the statistical power and diversity required to delineate robust, generalizable immunogenicity patterns and to advance toward a systems-level understanding of vaccine response.

However, translating such multifaceted data into actionable insights presents a new set of analytical challenges. Predictive methods must account for high interindividual variability, identify meaningful response patterns from limited samples, and incorporate baseline immune features to contextualize subsequent vaccine responses^10–14^. Emerging analytical tools have begun integrating diverse data types (e.g., transcriptomics, proteomics)^15–18^ but none has yet achieved a comprehensive, predictive framework that can navigate this complexity, accommodate outliers with atypical responses, and reliably anticipate who will benefit most from vaccination.

Achieving this predictive capability is essential for moving toward personalized vaccination strategies. Identifying immune markers that predict response heterogeneity can guide tailored interventions, ensuring that individuals less likely to respond robustly receive additional support. Such precision vaccinology approaches hold promise for enhancing vaccine effectiveness and coverage, especially in populations at higher risk^10^. Moreover, the ability to foresee vaccine responses is crucial for rapid and effective deployment of immunizations against pathogens with pandemic potential, including influenza virus and coronaviruses^19, 20^.

To meet these challenges, we introduce *immunaut*, an open-source, data-driven framework implemented as an R package for all systems vaccinologists aiming to unravel complex immune responses and predict vaccine outcomes. Through advanced dimensionality reduction, clustering, and predictive modeling, *immunaut* integrates multidimensional immune features to classify individuals into distinct immunophenotypic responder profiles, systematically revealing how baseline immune characteristics shape subsequent vaccine responses. Applied here to one of the most comprehensive LAIV datasets, *immunaut* delineated responder groups exhibiting systemic humoral, mucosal, or T-cell-mediated biases and uncovered critical biomarkers associated with effective LAIV responses. Beyond this specific application, *immunaut* is readily adaptable to additional vaccines and larger immunological datasets, offering a robust foundation for precision vaccinology and enabling researchers worldwide to develop more effective, tailored immunization strategies.

## Methods

### Study participants

We constructed a comprehensive dataset by compiling data from multiple research projects that evaluated immune responses to the trivalent live attenuated influenza vaccine (LAIV, Nasovac-S, based on the A/Leningrad/134/17/57 master donor strain, which is in use in Russia) among children in The Gambia. The cohort comprised 244 children aged 24-59 months who received LAIV during 2017 and 2018 as part of an open-label, prospective, observational, phase 4 immunogenicity study nested within a larger randomized trial (ClinicalTrials.gov identifier: NCT02972957)^7^. Eligible participants were healthy children with no history of respiratory illness in the preceding 14 days and no prior influenza vaccination. Exclusion criteria included serious active medical conditions (e.g., chronic diseases, severe malnutrition, genetic disorders), known immunodeficiency, hypersensitivity to vaccine components, recent use of immunosuppressive therapies, and contraindications to LAIV administration. Following community sensitization, recruitment was conducted in Sukuta, a peri-urban area in The Gambia. Written informed consent was obtained from parents or guardians, and the study was approved by The Gambia Government, the UK Medical Research Council Joint Ethics Committee, and the Medicines Control Agency of The Gambia, adhering to the International Conference on Harmonisation Good Clinical Practice standards. Participants received the Northern hemisphere formulation of LAIV (Nasovac-S) corresponding to their year of enrollment. Participants recruited in 2017 received the 2016-2017 formulation, which included strains A/17/California/2009/38 (H1N1) pdm09-like, A/17/Hong Kong/2014/8296 (H3N2)-like, and B/Texas/02/2013-like (B/Victoria/2/87-like lineage). In 2018, participants received the 2017–2018 formulation, where the H1N1 component was updated to A/17/New York/15/5364 (H1N1)pdm09-like, while the H3N2 and B strains remained unchanged. Vaccine titers per dose were consistent with manufacturer specifications for each strain and season. Whole blood and serum were collected at baseline (day 0) and post-vaccination on day 21. Nasopharyngeal swabs were collected at days 0, 2 and 7 post-LAIV using flocked swabs (Copan FLOQSwabs™) and stored in RNAprotect Cell Reagent (QIAGEN) for viral shedding assessment and microbiome analyses. To evaluate mucosal antibody responses, oral fluid samples were collected at days 0 and 21 post-LAIV using Oracol Plus swabs (Malvern Medical Development). Whole blood samples were drawn for serum separation, flow cytometry, and transcriptomic analyses. All samples were stored at −70°C until processing.

### Datasets

The datasets encompass a wide array of immune parameters

#### Humoral immune responses

Haemagglutination inhibition (HAI) assays^21^ were performed according to standard protocols using vaccine strain-matched antigens to assess seroconversion, defined as a fourfold or greater increase in HAI titers to ≥1:40 from day 0 to day 21^7^. This allowed for the evaluation of antibody responses against all 3 LAIV-vaccine strains: A(H1N1) pdm09, A(H3N2), and B/Victoria/2/87-like lineage influenza virus strains. Influenza virus-specific IgA in oral fluids was quantified using a protein microarray with recombinant haemagglutinin and neuraminidase proteins and normalized to total IgA in the sample (measured by enzyme-linked immunosorbent assay (ELISA))^22^. A twofold increase in the proportion of influenza virus-specific IgA was considered a significant mucosal antibody response. An influenza virus protein microarray (IVPM) was performed to determine the cross-reactive binding of serum antibodies against a panel of HA proteins from various influenza virus strains, including both vaccine-matched, drifted, and historical variants^23^. Antibody-dependent cellular cytotoxicity (ADCC) activity was assessed using reporter cell lines expressing Fc gamma receptors in the presence of chimeric H6/1 HA protein (H6 head domain combined with an H1 stalk domain), measuring the ability of antibodies to bind to group 1 HA stalk and mediate effector cell functions. ELISA based on standardized protocols was used to measure IgG levels to serum NA from N1 and N2, serum group 1 and 2 stalk-specific IgG using chimeric HA constructs (cH6/1 and cH7/3; H7 head domain on top of an H3 stalk domain), secretory IgA in oral secretions to N1 NA and group 1 stalk.

#### Cellular immune responses

T-cell responses before and on day 21 after LAIV were measured by stimulating fresh whole blood with overlapping 15-18-mer peptide pools covering vaccine-matched haemagglutinin (H1, H3, and B/Victoria/2/87-like HA), nucleoprotein (NP), and matrix (M) proteins. Intracellular cytokine staining for interferon-gamma (IFN-γ) and interleukin-2 (IL-2) was performed, and responses were analyzed using flow cytometry, as previously described^7^.

#### Viral shedding, microbiome analysis, and viral load

Nasopharyngeal swabs were assessed for LAIV strain shedding on days 2 and 7 post-LAIV using reverse-transcription PCR (RT-PCR) assays targeting haemagglutinin genes as previously described^7^. Quantitative RT-PCR provided viral load measurements expressed as log₁₀ egg infectious dose equivalents per mL. Additionally, the presence and density of nasopharyngeal *Streptococcus pneumoniae* before vaccination were quantified as previously described^9^. Baseline samples were tested for the presence of respiratory viruses using a multiplex real-time PCR method, as detailed in the original publication^24^. The assay panel included influenza A and B viruses, respiratory syncytial virus (RSV) types A and B, human parainfluenza viruses (HPIV) 1-4, human metapneumovirus, adenovirus, seasonal coronaviruses (229E, OC43, NL63), and human rhinovirus.

#### Immunophenotyping

Multicolor flow cytometry panels were utilized to quantify frequencies of innate immune cell subsets before vaccination. The cell populations analyzed included myeloid dendritic cells (mDCs), plasmacytoid dendritic cells (pDCs), monocyte subsets (classical, intermediate, and non-classical monocytes), and T follicular helper (Tfh) Cells: Circulating Tfh cells expressing activation markers (CXCR3⁺ICOS⁺PD-1⁺) were quantified at baseline to assess their role in supporting antibody responses^24^.

#### Transcriptomic profiles

RNA sequencing was conducted on nasal swabs collected before LAIV to generate transcriptomic profiles^8^. Gene set enrichment analysis (GSEA) was used to identify gene expression signatures and pathways associated with vaccine response and immune modulation.

#### Demographic and clinical data

Detailed demographic data, including age, sex, nutritional status (e.g., weight-for-height Z-score), and health history, were collected to assess potential correlations with immune responses. Participants were monitored for adverse events, and any respiratory illnesses occurring during the study period were documented to evaluate safety and potential confounding factors.

### Data integration and preprocessing

The integrated dataset was generated using the standard extract-transform-load (ETL) procedure, as described previously^17^. Briefly, data from six primary datasets, each provided in CSV format and encompassing various immunological assays and demographic information, were integrated using the unique identifier ‘*Subject ID’*. This integration was facilitated by a custom ‘*combine_data’* function, which merged the datasets into a single comprehensive dataset. Data were obtained at baseline (day 0) and day 21 post-vaccination for all measured parameters, including cellular, humoral, and mucosal values. Fold-changes were then calculated to obtain the LAIV-responsiveness measures, capturing both the pre-existing immune state and the vaccine-induced responses. Before analysis, the integrated dataset underwent several preprocessing steps. Missing values were imputed using median imputation (*medianImpute*), where missing entries were replaced with the median value of the respective feature. The data were then normalized by centering (subtracting the mean) and scaling (dividing by the standard deviation) of each feature. Features exhibiting zero variance (*zv*) and near-zero variance (*nzv*) were identified and removed to reduce noise and improve computational efficiency. Additionally, features with pairwise Pearson correlation coefficients greater than 0.85 were considered highly correlated and were removed to mitigate multicollinearity issues. The final dataset included a comprehensive set of immunological and demographic features representing various aspects of the immune response to LAIV.

### Data-driven immunogenicity responders subtyping

In this section, we describe the methodology used for clustering a dataset based on t-SNE dimensionality reduction^25^, K-Nearest Neighbors (KNN) graph construction, and Louvain community detection^26, 27^. We also outline the optimization steps for selecting the best clustering result based on multiple clustering evaluation metrics.

### t-SNE dimensionality reduction

Let **X** *∈ Rn×d* represent the dataset with *n* samples and *d* features. We first apply t-SNE to project the dataset into a lower-dimensional space **Y** *∈ Rn×*2. The t-SNE method aims to minimize the Kullback-Leibler (KL) divergence between probability distributions of points in high-dimensional and low-dimensional spaces. The objective function minimized by t-SNE is:

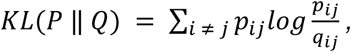

where *p_ij_*is the similarity between points *i* and *j* in the high-dimensional space, and *q_ij_* is the similarity in the low-dimensional space.

### K-nearest neighbors (KNN) graph construction

Given the t-SNE projection Y, we construct a K-Nearest Neighbors (KNN) graph to capture the local structure of the data. For each point *i*, the *k* nearest neighbors are determined based on the Euclidean distance in the 2D space:

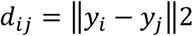

where y*i* and y*j* are the t-SNE coordinates of points *i* and *j*, respectively. The graph *G* = (*V, E*) is constructed with *V* being the set of nodes (samples) and *E* the set of edges connecting each point to its *k* nearest neighbors. The weight of each edge is defined as:

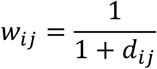

where smaller distances lead to higher edge weights, emphasizing closer neighbors.

### Louvain clustering for community detection

The Louvain method is applied to the KNN graph for community detection. The Louvain algorithm optimizes *modularity Q*, which measures the density of edges within communities compared to what would be expected in a random graph. The modularity is defined as:

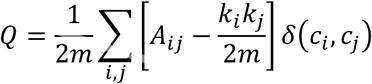

where:

- *Aij* is the adjacency matrix of the graph,
- *ki* is the degree of node *i*,
- *m* is the total number of edges,
- *ci* is the community assignment of node *i*,
- *δ*(*ci, cj*) is the Kronecker delta function that equals 1 if *ci* = *cj* and 0 otherwise.

The Louvain method iteratively maximizes *Q* by merging nodes and communities to achieve an optimal partitioning.

### Iterative optimization of clustering resolution

To explore different clustering resolutions, we apply the Louvain algorithm over a range of resolutions *r*. The resolution *r* controls the granularity of the clustering, with lower resolutions favoring fewer, larger clusters, and higher resolutions producing more, smaller clusters. We define a sequence of resolutions {*r*_1_, *r*_2_, …, *r*_*k*_} such that: *r*_*i*+1_ = *r*_*i*_ + Δ*r*, Δ*r* = 0.1 for each iteration *i*. For each resolution *r_i_*, we compute the modularity *Q*(*r_i_*) and the number of clusters *C*(*r_i_*). We keep the clustering results that fall within the desired range of cluster counts: *C*_min_ *≤ C*(*r_i_*) *≤ C*_max_.

### Evaluation metrics for best clustering selection

Once we obtain multiple clustering results across different resolutions, we select the best result based on a combination of metrics:

#### Modularity Q

We aim to maximize the modularity score, which indicates better separation of communities.

#### Silhouette score S

The silhouette score measures the cohesion and separation of clusters. For each point *i*, the silhouette score is defined as:

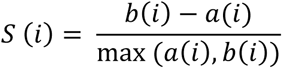

where *a*(*i*) is the average distance between *i* and all other points in the same cluster, and *b*(*i*) is the average distance between *ii* and all points in the nearest cluster. We maximize the average silhouette score across all points.

#### Davies-Bouldin index (DBI)

The DBI is computed as:

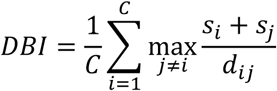

where *s*_*i*_ is the average distance within cluster *i*, and *d_ij_* is the distance between cluster centroids *i* and *j*. A lower DBI indicates better clustering.

*Calinski-Harabasz index (CH):* The CH index is given by:

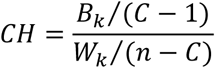

where *B*_*k*_ is the between-cluster dispersion and *W*_*k*_ is the within-cluster dispersion. Higher CH values indicate better clustering.

### Combined score for clustering selection

For each clustering result, we normalize the metrics and compute a combined score *M* to select the best clustering: *M* = *α*1 *·* normalize(*Q*) + *α*2 *·* normalize(*S*) + *α*3 *·* (1 *-* normalize(*DBI*)) + *α*4 *·* normalize(*CH*), where *α*1*, α*2*, α*3*, α*4 are weights assigned to each metric, and the normalization function scales each metric to the range [0, 1]. The clustering result with the highest score, *M,* is selected as the final optimal clustering.

### Predictive modeling of immunophenotypic clusters

Following unsupervised ML clustering, the immunophenotypic groups identified in *immunaut*’s first step were treated as categorical outcomes in a predictive modeling framework. In this second step, the Sequential Iterative Modeling OverNight (SIMON) platform^17, 28^ was employed to systematically evaluate 141 machine learning (ML) algorithms, aiming to discover a minimal set of baseline features capable of accurately predicting immunophenotypic group membership. Predictors were baseline measurements of immune and molecular features, with immunophenotypic groups from clustering serving as the outcome variable. Data preprocessing procedures included centering and scaling, median imputation for missing values, removal of highly correlated features, and zero- and near-zero-variance filtering to ensure data quality. The dataset was divided into 80% training and 20% testing sets for model development, allowing for independent model validation. Parallel computation was implemented to expedite the training and selection process, with the number of cores for parallel processing set to the number of available CPU cores minus one. Model evaluation during training utilized a 10-fold cross-validation approach, repeated three times to enhance robustness and mitigate overfitting. The performance of each model was assessed on the independent test set, using a confusion matrix and area under the curve (AUC) metrics to provide unbiased evaluations of predictive accuracy across the three response classes. One-vs-all receiver operating characteristic (ROC) curves were generated for each class using the *pROC* package in R, allowing for a detailed assessment of model sensitivity and specificity. To gain insights into feature significance, variable importance scores were calculated for each model within each response class. These scores were aggregated across classes to highlight baseline features with the highest predictive power, providing a comprehensive view of the immune and molecular markers most strongly associated with specific immunophenotypic group memberships.

### Model interpretability

*<u>SHAP (SHapley Additive exPlanations) analysis</u>* was conducted using the DALEX (moDel Agnostic Language for Exploration and eXplanation) package in R (https://github.com/ModelOriented/DALEX/) to interpret the contribution of individual features to the gbm model’s predictions for each LAIV responder group^29, 30^. SHAP values were computed to quantify the local, observation-specific impact of each feature on the model’s output, providing an additive decomposition of predictions into contributions from individual features and an intercept term. For each observation, SHAP values reflect how much each feature increases or decreases the predicted probability of belonging to a specific cluster (Group 1: CD8 T-cell responders, Group 2: mucosal responders, Group 3: systemic, broad influenza A responders) relative to the baseline prediction (intercept). The analysis was implemented by linking the trained gbm model with the DALEX explainer function, generating SHAP values for features prioritized by global variable importance scores. Feature contributions were visualized for each cluster using horizontal bar plots, where the magnitude and direction of SHAP values indicate the relative importance and influence (positive or negative) of each feature on the prediction. This approach provided granular insights into how baseline immune features drive LAIV immunogenicity across different responder groups. *<u>Tree-based analysis.</u>* All analyses were conducted in R using the *rpart* and *rpart.plot* packages. Data were loaded from a CSV file and merged with feature mapping information to restore original feature names. Missing values were replaced by column medians to ensure complete datasets for model fitting. Categorical variables were converted to factors, and continuous variables were discretized into meaningful bins based on predefined cutoffs. After discarding redundant variables, a decision tree model was fitted using *rpart* with parameters set to ensure appropriate pruning (cp=0.01) and sufficient sample sizes for splits (minsplit=70, minbucket=10). The tree was visualized with *rpart.plot*, and its full rule set was extracted using *rpart.rules* and saved for downstream interpretation.

### Data analysis

Statistical analysis was performed using R (https://www.r-project.org/) package *ggpubr* version 0.4.0. Integrative and machine learning analysis, including hierarchical clustering, t-SNE, KNN, and Louvain clustering, and supervised ML approach SIMON, were performed using PANDORA software version 0.2.1. All data visualizations were conducted in R version 4.3.1 with the *tidyverse* package (version 2.0.0) for data wrangling. Heatmaps were created using the *pheatmap* package (version 1.0.12), polar plots were produced with *ggplot2* (version 3.5.1) and the *Wes Anderson color palette* (version 0.3.7), and radar plots were generated with *fmsb* (version 0.7.6). Scaled median pathway expression values were calculated by grouping genes by pathway, omitting any missing values, and computing the median for each pathway-group pair. These scaled median values were used in all visualization techniques for consistent metric comparison across clusters in each plot type. Feature-specific polar plot values were further transformed using log10 to control significant variances, ensuring a more balanced visualization of expression levels across features.

### Data availability

The complete integrated dataset supporting the findings in this study is available on Zenodo^31^: <u>Comprehensive Multimodal Immune Response Dataset for LAIV</u> <u>Vaccination in Pediatric Cohorts</u>. This dataset includes all baseline and post-vaccination measurements required to reproduce the analyses presented in this study. The *immunaut* platform, used for mapping immune profiles and predicting vaccine responses, is accessible via the PANDORA AI platform (https://pandora.atomic-lab.org/) and as an R package on CRAN (https://cran.r-project.org/web/packages/immunaut/index.html). Detailed documentation for *immunaut* is provided on the GitHub repository (https://github.com/atomiclaboratory/immunaut).

## Results

### Comprehensive immunoprofiling of LAIV responses reveals distinct immunophenotypic groups

To define responder status to LAIV in a cohort of 244 Gambian children^7^, we focused on adaptive immune markers (systemic and mucosal) for which we had paired baseline (day 0) and post-vaccination (day 21) measurements and expressed these as fold-change values (V21/V0; see **Materials and Methods**) (**Fig. 1A**). These included key humoral and cellular immune parameters, enabling us to quantify LAIV-induced immunogenicity relative to each participant’s pre-vaccination baseline. By using fold-change rather than absolute values, we accounted for interindividual variability in baseline immunity, ensuring that our clustering captured genuine vaccine-induced changes. Within this framework, we evaluated a comprehensive panel of antibody-mediated responses. We measured hemagglutination inhibition (HAI) titers, an indicator of antibodies that block the binding of the influenza virus to host cells^21^. We also employed the influenza virus protein microarray (IVPM) to assess the breadth of antibody responses elicited by vaccination^32^. The IVPM is a high-throughput hemagglutinin (HA) microarray platform designed to profile binding antibody responses across multiple influenza virus strains. This HA-based microarray includes HA proteins from a diverse panel of influenza A and B viruses, enabling the simultaneous assessment of antibody reactivity to various subtypes and lineages. In our analysis, we incorporated HA antigens from influenza A H1N1 virus strains such as A/California/07/2009 (CAL09 H1 IVPM) and A/Michigan/45/2015 (MICH15 H1 IVPM), as well as H1 isolates A/New Caledonia/20/1999 (NC99 H1 IVPM) and A/Guangdong Maonan/SWL1536/2019 (GD H1 IVPM). For influenza A virus H3N2, we included HA proteins from A/Switzerland/9715293/2013 (SWISS H3 IVPM), A/Hong Kong/4801/2014 (HK14 H3 IVPM), and A/Kansas/14/2017 (KAN H3 IVPM). Additionally, the microarray featured HA antigens from influenza B viruses, specifically B/Phuket/3073/2013 (B/Yamagata/16/88-like lineage) and B/Washington/02/2019 (B/Victoria/2/87-like lineage). By utilizing the IVPM with this comprehensive set of HA antigens, we quantitatively evaluated serum antibody binding profiles before and after LAIV administration. The HA microarray allows us to obtain detailed insights into the specificity, magnitude, and breadth of the antibody responses elicited by the vaccine. Specifically, it enables the detection of antibodies that bind to a wide range of HA antigens, including those from strains not present in the vaccine formulation. This allows us to assess the extent of cross-reactive antibody responses, which is crucial for understanding the potential for broad protection against diverse influenza viruses. Additionally, we examined stalk-specific responses targeting conserved regions of the hemagglutinin (HA) protein, including antibody-dependent cellular cytotoxicity (ADCC) activity measured against chimeric HA stalk constructs like cH6/1 and cH7/3, which are engineered to measure antibodies to the stalk domain, enabling assessment of cross-reactive immunity to influenza virus strains beyond strain-specific head responses^33^. Neuraminidase (NA) titers were also assessed in blood and nasal mucosa, offering insights into cross-protective responses across this surface protein^34^. Complementing the antibody profiles, we assessed T-cell interferon-gamma (IFN-γ) and interleukin-2 (IL-2) production upon stimulation with vaccine strain components such as hemagglutinin (HA), neuraminidase (NA), and the matrix/nucleoprotein (M/NP) proteins, capturing the magnitude and diversity of systemic cellular responses.

**Figure 1.**
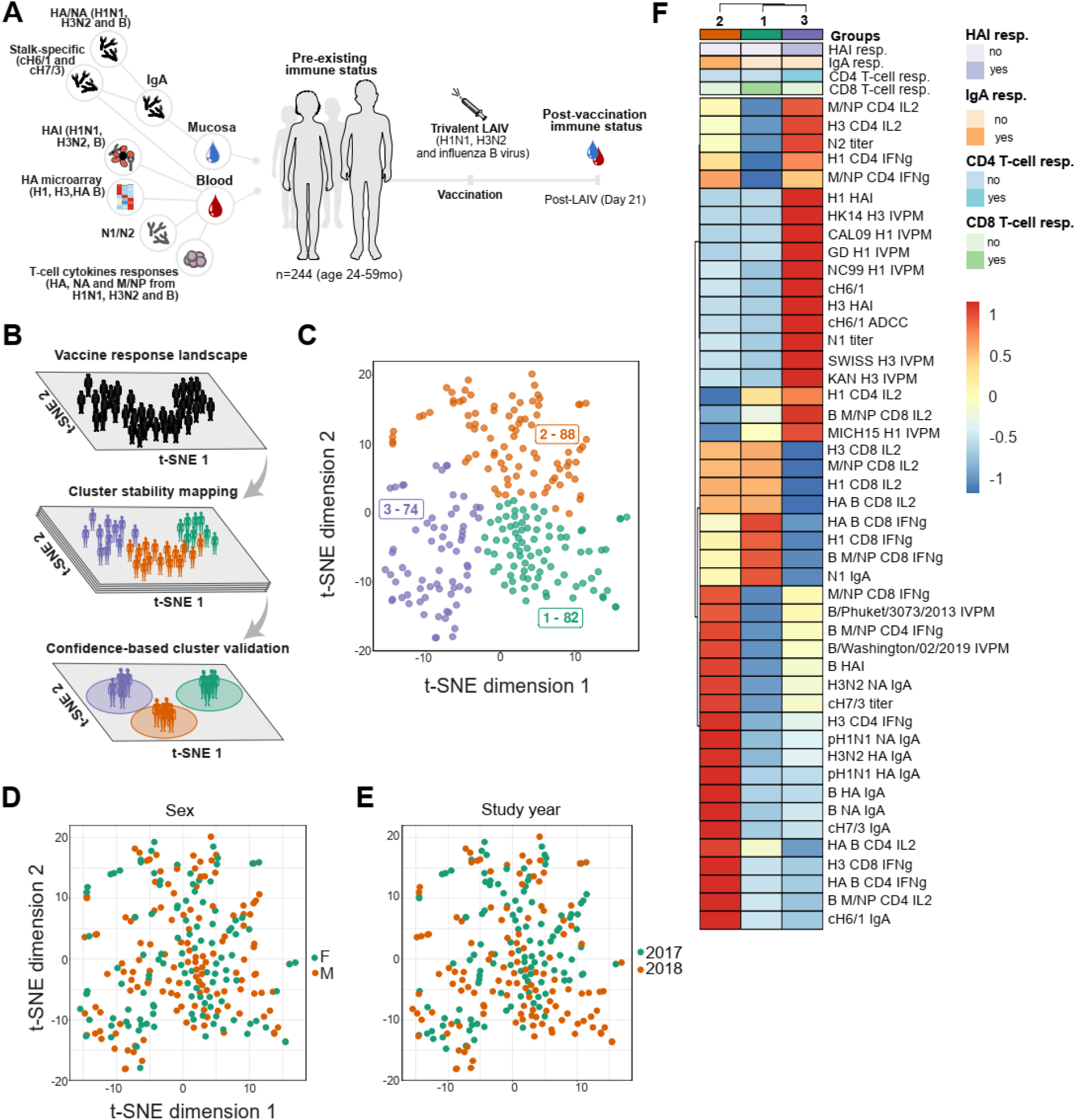
Immune response landscape mapping of LAIV reveals distinct immunophenotypic groups. **(A)** Cohort overview depicting all features that were used for unsupervised ML approach: 244 children (24-59 months) vaccinated with LAIV, with mucosal and blood samples collected on day 0 (baseline) and day 21 (post-vaccination) to capture vaccine-induced immune responses by accounting for pre-existing immune status (fold-change). **(B)** Workflow schematic for automated clustering pipeline for immune landscape generation, applying t-SNE dimensionality reduction, K-nearest neighbors (KNN) graph construction, and Louvain community detection to identify distinct immunophenotypic clusters. The dataset is first reduced to a two-dimensional space using t-SNE to retain local similarities while capturing high-dimensional data structure generating a vaccine response landscape for each individual. A KNN graph is then constructed to capture local data relationships in this reduced space. Louvain clustering is applied to the KNN graph, optimizing community modularity by maximizing intra-cluster density. To ensure map stability, multiple clustering resolutions are iteratively tested and evaluated using a combined score based on modularity, silhouette score, Davies-Bouldin index (DBI), and Calinski-Harabasz index (CH), selecting the best clustering configuration based on these metrics. **(C)** Clustered t-SNE plot of fold-change data (post/pre-LAIV) revealing three LAIV response phenotypes (perplexity: 30; exaggeration factor: 4; max iterations: 10,000; theta: 0; eta: 500; K: 60; silhouette score: 0.40). **(D, E)** Each panel highlights clustering patterns according to specific demographic and response factors; **(D)** Clustering by sex (male in orange and female in green) **(E)** Clustering by study year (2017 in green and 2018 in orange). **(F)** Heatmap and hierarchical clustering display fold-change (FC) data for each cluster, scaled from −1 to 1. Clustering uses Euclidean distance and Ward’s D2 method, with cluster ordering optimized for visual clarity.

Collectively, this panel of immunological assays provided a highly granular view of the magnitude and quality of immune responses elicited by LAIV, allowing us to capture a detailed immunophenotypic landscape that includes systemic and mucosal humoral and cellular responses, cross-reactive and subtype-specific functional activity, as well as neutralization potential, thereby offering robust insights into the breadth and functional diversity of vaccine-induced immunity.

This integrated, multimodal dataset containing the abovementioned fold-change parameters served as input for *immunaut* unsupervised machine learning framework for exploring the vaccine response landscape (**Fig. 1B**, for a full technical description, see the **Materials and Methods**). Using t-distributed stochastic neighbor embedding (t-SNE) for dimensionality reduction, we projected the high-dimensional immune response data into a lower-dimensional space to reveal patterns and relationships across individuals. Following this, we constructed a weighted graph where nodes represent individuals, and edges reflect proximity based on Euclidean distances in the reduced space, with higher weights assigned to closer neighbors. Louvain clustering was then used to maximize modularity, iteratively grouping samples into distinct immunophenotypic clusters. By iteratively refining cluster granularity and assessing cluster stability, we established robust immunophenotypic clusters, effectively capturing the variability of immune responses to LAIV within this diverse cohort (**Fig. 1C**). This unbiased analysis generated three distinct groups with shared immune response profiles (**Fig. 1C**). Specifically, Group 1 (green) contains 82 individuals, Group 2 (orange) includes 88 individuals, and Group 3 (purple) comprises 74 individuals.

To ensure that the immunophenotypic clusters identified in our analysis accurately reflect distinct immune responses to LAIV, we hypothesized that the clustering should be robust and not merely a reflection of demographic variables or external biases. To test this, we first assessed the quality of cluster separation using the silhouette score, which yielded a value of 0.4. This score indicates a moderate level of cluster distinction, supporting the validity of our clustering while accounting for the expected biological variability inherent in immune responses among individuals. To further validate our approach, we investigated whether factors not included in the immunological analysis - specifically demographic variables such as gender and study year -influenced the clustering outcome (**Fig. 1D, E**). Gender differences are known to affect immune responses to vaccines^35–37^, and variations in vaccine strains across study years (e.g., different H1N1 strains included in the vaccine formulations) could potentially impact the immune profiles^7^. Upon examining the distribution of these variables across the clusters, we observed minimal overlap, suggesting that the clusters are not driven by demographic factors (**Fig. 1D**) or study year differences (**Fig. 1E**). This demonstrates that the clustering captures three distinct immunophenotypic landscapes induced by LAIV vaccination rather than reflecting predefined biases or demographic influences. This conclusion supports the robustness of our method and confirms that the identified clusters represent genuine differences in immune responses to LAIV.

In our analysis, individuals in Group 1 (n=82) demonstrated a distinct immunological profile characterized predominantly by CD8 T-cell-mediated responses following LAIV vaccination (**Fig. 1F**). Specifically, this group exhibited increases in CD8 T-cell activity, as evidenced by elevated interferon-gamma (IFN-γ) and interleukin-2 (IL-2) production upon stimulation with influenza virus antigens (**Fig. 1F**). The most pronounced responses were observed against HA and M/NP antigens from influenza B virus (**Fig. 1F**). Notably, the induction of HA B-specific CD8 T-cell IFN-γ responses reached statistical significance when compared to Group 3, indicating a robust cellular immune activation targeting this viral component (**Fig. 2A**). Conversely, humoral responses in Group 1 were minimal or absent (**Fig. 1F**; **Fig. 2A, D**). There was a lack of increases in HAI titers, and antibody binding responses assessed via the HA microarray were negligible, suggesting that these individuals did not mount a substantial antibody-mediated response to the vaccine (**Fig. 1F**; **Fig. 2A, D**). IgA responses were also lacking in this group. Although some N1-specific IgA responses were detected (**Fig. 1F**), these were not statistically significant and were comparable to those observed in Group 2, indicating that the N1 IgA responses were not a distinguishing feature of Group 1 (**Fig. S1A**). Based on these results, we term individuals in Group 1 as ‘*CD8 T-cell responders*.’ These findings suggest that, in Group 1, LAIV vaccination induces a cellular immune response characterized by enhanced CD8 T-cell activity against specific influenza antigens, particularly those associated with the influenza B virus.

**Figure 2.**
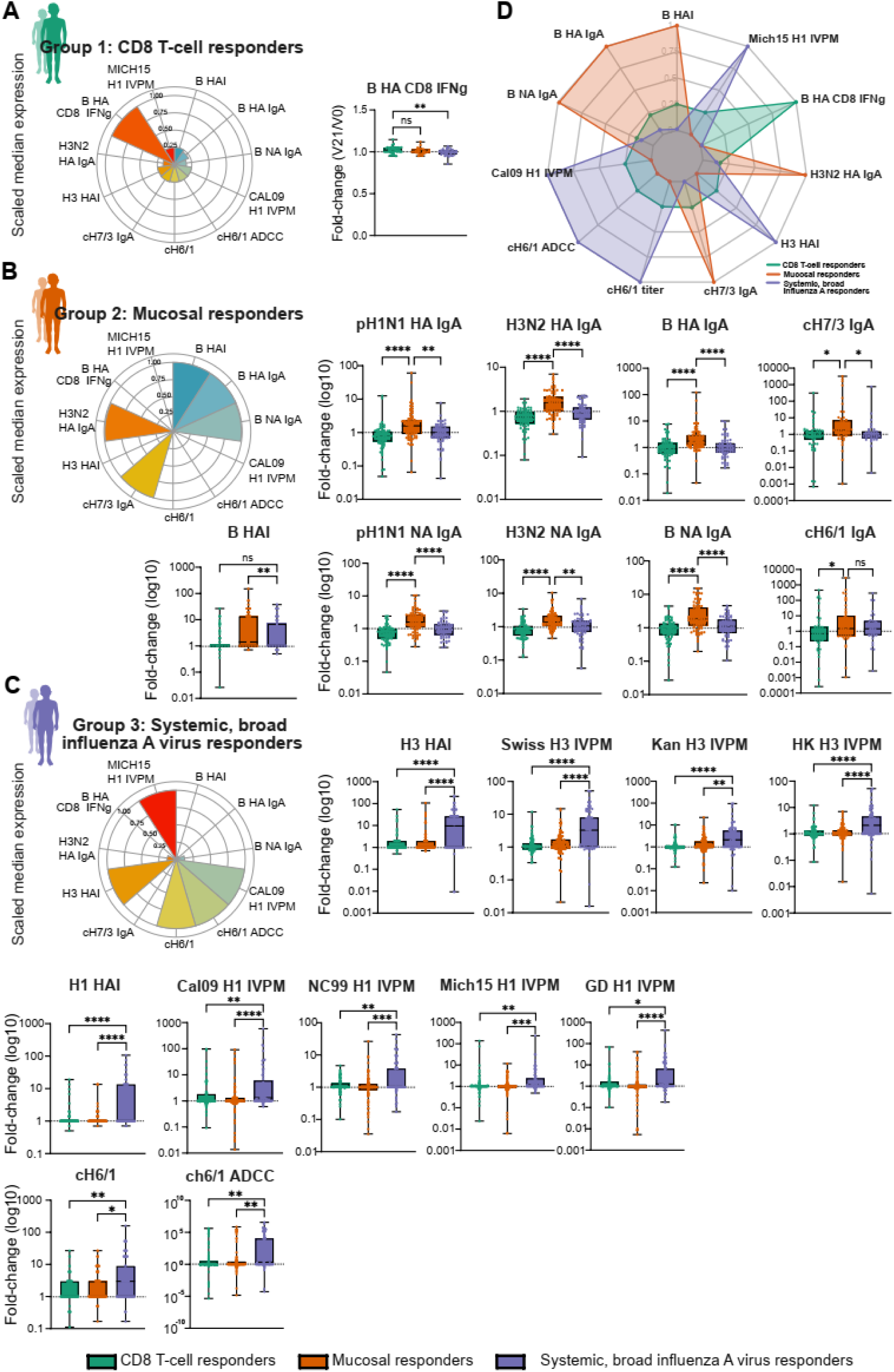
Vaccine response immune signatures defining LAIV responder types. **(A)** Polar plot summarizing scaled median expression of key immune features in CD8 T-cell responders (Group 1, green). CD8 T-cell responders are characterized by robust influenza B virus HA-specific CD8+ IFNγ responses and limited humoral immunity, with median feature values represented in the polar plot and fold-change comparisons shown in the adjacent box plot. **(B)** Polar plot for mucosal responders (Group 2, orange), illustrating strong mucosal IgA responses, particularly stalk-specific (cH7/3 IgA) and H3N2 virus HA-specific IgA antibodies and influenza B virus-specific responses. Box plots detail fold changes (shown as log10) for various immune features, highlighting systemic (influenza B virus HAI) and mucosal immune activation (IgA). **(C)** Polar plot depicting systemic, broad influenza A virus responders (Group 3, purple), showing elevated systemic antibody responses to multiple influenza A virus strains (e.g., H1, H3), as well as cross-reactive IgG and ADCC (antibody-dependent cellular cytotoxicity) activity. Box plots show fold-change values (log10) for each immune marker across responder groups. **(D)** Integrated radar plot comparing scaled median immune expression profiles across all responder groups (CD8 T-cell responders in green, mucosal responders in orange, systemic broad influenza A virus responders in purple), emphasizing distinct immune feature distributions. This integrative visualization highlights the unique baseline and post-vaccination immune landscapes that define each responder profile. Box plots denote min to max values, points are all individuals within the group, with significance levels calculated using one-way ANOVA Kruskal-Wallis test with Dunn’s multiple comparison test to adjust for multiple testing. Significance is indicated as follows: ns = not significant, *p < 0.05, **p < 0.01, ***p < 0.001, ****p < 0.0001.

In contrast to Group 1, Group 2 (n=88) individuals exhibited an immunological profile predominantly characterized by mucosal IgA responses following LAIV vaccination (**Fig. 1F**). This group demonstrated statistically significant induction of mucosal IgA antibodies across all antigens and strains tested, including pandemic H1N1 (pH1N1) HA and NA, H3N2 HA and NA, and B/Victoria/2/87-like lineage HA and NA (**Fig. 2B**). Elevated secretory IgA responses were also observed against chimeric hemagglutinin group 1 (cH6/1) and group 2 (cH7/3) stalk constructs (**Fig. 2B**), indicating the induction of antibodies targeting conserved regions of the HA protein. The consistent and statistically significant increase in mucosal IgA responses was unique to Group 2 and not observed in Groups 1 or 3 (**Fig. 2B, D**), validating our classification of these individuals as ‘*Mucosal responders*.’ In addition to mucosal IgA responses, Group 2 also exhibited significant seroconversion to influenza B viruses (**Fig. 1F**). This was evidenced by statistically significant increases in HAI titers against influenza B virus (**Fig. 2B**). Although antibody binding responses to B/Phuket/3073/2013 (B/Yamagata/16/88-like lineage) and B/Washington/02/2019 (B/Victoria/2/87-like lineage) measured by the HA microarray were elevated in Group 2 (**Fig. 1F**), these increases did not reach statistical significance (**Fig. S1B**). The enhanced seroconversion to the influenza B virus indicates that the humoral immunity in Group 2 extends beyond mucosal surfaces to include systemic antibody responses against the influenza B virus.

In contrast to Groups 1 and 2, Group 3 (n=74) individuals exhibited an immunological profile predominantly characterized by robust systemic antibody responses to influenza A viruses following LAIV vaccination (**Fig. 1F**). This was evidenced by significant increases in HAI titers for both H1N1 and H3N2 strains (**Fig. 2C**), indicating effective seroconversion to these viruses. Further analysis revealed that the antibody responses in Group 3 extended beyond the vaccine strains, demonstrating a breadth of reactivity against multiple influenza A viruses. Significant increases in antibody binding responses were observed to HA subtypes from H1N1 strains - A/California/07/2009 (Cal09 H1 IVPM), A/Michigan/45/2015 (Mich15 H1 IVPM), A/Guangdong Maonan/SWL1536/2019 (GD H1 IVPM), and A/New Caledonia/20/1999 (NC99 H1 IVPM) - and H3N2 strains - A/Switzerland/9715293/2013 (SWISS H3 IVPM), A/Kansas/14/2017 (KAN H3 IVPM), and A/Hong Kong/4801/2014 (HK14 H3 IVPM) (**Fig. 2C**). This antibody breadth includes historical and drifted strains not present in the vaccine formulation, suggesting the induction of cross-reactive immunity capable of recognizing contemporary and historical influenza A viruses. The ability of these children, born between 2012 and 2015, to mount such broad antibody responses suggests that LAIV can effectively stimulate immunity against both current and historical influenza A strains, enhancing the potential for cross-protection against antigenic drift. Elevated responses were also observed against the chimeric hemagglutinin construct cH6/1, including increased antibody-dependent cellular cytotoxicity (ADCC) activity targeting cH6/1 (**Fig. 2C**). This indicates the induction of antibodies against conserved HA stalk regions of group 1 influenza A viruses. Additionally, significantly higher N1 titers were detected in Group 3 (**Fig. S1C**), further supporting a coordinated immune response targeting conserved epitopes in both the HA stalk and NA proteins associated with H1N1 viruses. While CD4 T-cell responses producing IFN-γ and IL-2 in response to M/NP peptides from influenza A viruses were significantly elevated in this group, increases in H3 CD4 T-cell IL-2 and H1 CD4 T-cell IFN-γ were observed at low levels (**Fig. S1C**). Notably, the absence of significant IgA responses in Group 3 indicates that immunity in these individuals is predominantly systemic rather than mucosal (**Fig. 2B, D**). This lack of mucosal antibody production, combined with robust systemic antibody responses and cross-reactive immunity against influenza A viruses, supports classifying these individuals as ‘*Systemic broad influenza A virus responders*.’ Their immunoprofile demonstrates that LAIV vaccination in this group elicits a potent, cross-reactive systemic antibody response targeting multiple influenza A viruses, including both H1N1 and H3N2 strains. This emphasizes the vaccine’s capacity to induce protective immunity against various influenza A viruses.

Our comprehensive integrative analysis reveals these nuanced immunoprofiles by considering multiple immunological parameters simultaneously. Unlike traditional approaches focusing on a single immune response (such as HAI titers), our method integrates data across humoral and cellular responses at systemic and mucosal levels, providing deeper insights into the immune landscape elicited by LAIV vaccination. This holistic approach highlights the superiority of our methodology in capturing the key protective mechanisms - specifically, the robust systemic antibody responses against a diverse array of influenza A viruses observed in Group 3 - that would otherwise remain obscured.

### Predictive modeling of LAIV response phenotypes based on baseline immune profiles

We next sought to determine if pre-vaccination immune profiles could predict an individual’s likelihood of responding to LAIV as a CD8 T-cell, mucosal, or systemic, broad influenza A virus responder. To achieve this, we used comprehensive baseline immunological measurements capturing various immune parameters before vaccination (**Fig. 3A**). These included baseline measurements of antibody profiles (HAI titers, IVPM assays), stalk-specific antibody responses (cH6/1 and cH7/3 constructs and ADCC measurements) and T-cell responses (ICS assays across HA, NA, and M/NP proteins for influenza A and B viruses). We also included baseline *S. pneumoniae* load (pneumococcal carriage) and asymptomatic respiratory viral presence to account for potential modulators of mucosal immunity, as these factors can shape immune readiness and impact LAIV response patterns^8, 38^. Baseline RNA pathway scores from single-sample Gene Set Enrichment Analysis (ssGSEA) on nasal samples provided insights into activities in innate immunity, cytokine signaling, chemokine activity, and cellular defense pathways^8^. Additionally, the baseline frequency of immune cell subsets such as classical, intermediate, and non-classical monocytes, plasmacytoid and myeloid dendritic cells (pDCs/mDCs), and T follicular helper (TFH) cells^24^ were evaluated due to their roles in immune activation and regulation relevant to LAIV, which relies on both innate and adaptive immune pathways to induce protection^5, 6, 39, 40^.

**Figure 3.**
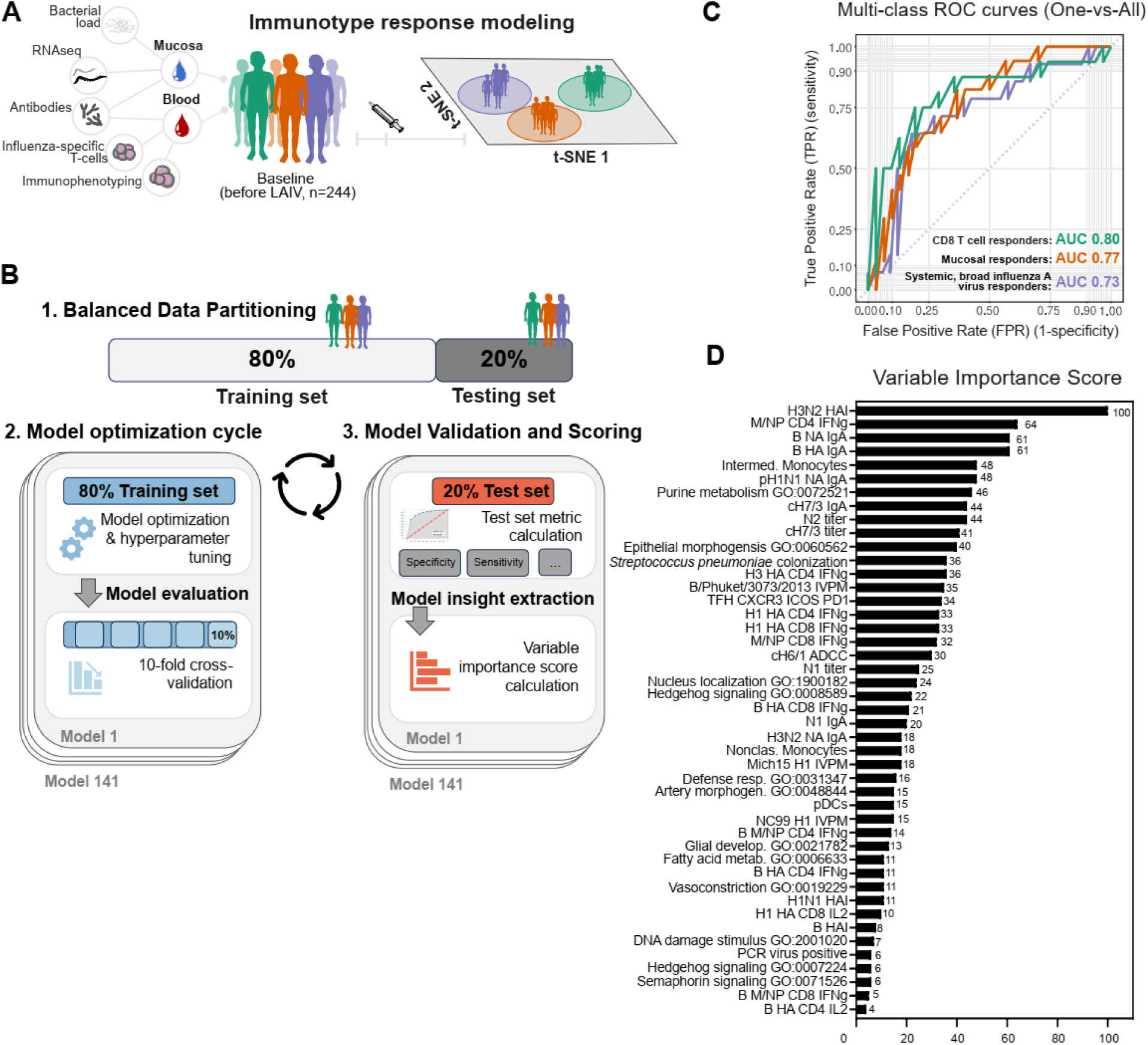
Automated machine learning framework for mapping and predicting LAIV immunogenicity response phenotypes. **(A)** Overview of the automated ML framework developed to predict LAIV response phenotypes using baseline immune data from mucosal and blood samples, capturing multi-dimensional immune parameters such as transcriptomics, antibody titers, bacterial load, flu-specific T-cell responses, and comprehensive immunophenotyping. **(B)** *Step 1.* Balanced data partitioning: the dataset is split into training (80%) and testing (20%) sets, ensuring proportional representation of each immunophenotypic group (CD8 T-cell, mucosal, and systemic, broad influenza A responders) to maintain predictive accuracy across classes. *Step 2.* Model optimization cycle: 10-fold cross-validation and hyperparameter tuning are applied across 141 machine learning models, each iteratively trained and validated to identify the best predictors of vaccine response. *Step 3.* Model evaluation and scoring: predictive performance metrics, including specificity, sensitivity, and area under the curve (AUC), are calculated on the test set (20%) for model validation. Feature importance scores are computed for each baseline variable, providing a ranked analysis of each immune parameter’s contribution to LAIV response prediction. **(C)** Multi-class ROC plot of the gbm model evaluated on the test set (20%), displaying predictive accuracy across all three classes: CD8 T-cell responders (green), mucosal responders (orange), and systemic, broad influenza A responders (purple) in a one-vs-all comparison. **(D)** Variable importance score table for the gbm model, showcasing the cumulative importance of the selected baseline features across the three predicted classes, highlighting the most influential parameters in LAIV immunogenicity prediction.

To model the mapped vaccine responses and identify baseline predictors of LAIV response types, we applied the Sequential Iterative Modeling OverNight (SIMON) platform^17, 28^ (**Fig. 3B**). SIMON is specifically designed to handle high-dimensional datasets with diverse immune parameters and substantial interindividual variability^17^, making it highly suitable for our complex dataset. Our methodological approach involved systematically testing 141 ML algorithms using the SIMON platform to explore various modeling techniques. This comprehensive screening is important because different algorithms excel at capturing various forms of nonlinear patterns, interactions, and dependencies within high-dimensional biological data^17, 28, 41, 42^. By encompassing ensemble methods, regularization techniques, and nonlinear classifiers, we aimed to ensure the selection of the most accurate and biologically meaningful model for our dataset. We employed rigorous 10-fold cross-validation during model training to enhance robustness and mitigate overfitting, and we assessed performance on a held-out test set to ensure generalizability. Out of the 141 models tested, 77 were successfully trained and evaluated, with 26 achieving a test set area under the receiver operating characteristic curve (AUC) above 0.7 (**Supplementary Table S1**). Models exceeding an AUC of 0.7 demonstrated a strong capacity to distinguish among response classes, underscoring the predictive strength of our baseline immune profiles.

Among all models evaluated, the gradient boosting machine (gbm) model emerged as the top-performing algorithm, evaluated stringently across multiple performance measures, confirming its suitability for this multiclass classification task (**Supplementary Table 1**). In addition to the overall and class-specific metrics derived from the confusion matrix on the held-out test set (20% of the data), the gbm model was selected based on its comprehensive performance profile. The gbm model achieved an accuracy of 59.57%, substantially exceeding the null accuracy of 36.17% (p = 0.0009), and demonstrated a balanced accuracy of 71.67%, reflecting its ability to perform well across all classes. Its Kappa statistic (0.3929) indicates fair agreement beyond chance, while the F1-score (0.6286), precision (0.6902), and recall (0.6471) highlight its capacity to balance false positives and false negatives. The high negative predictive value (0.8065) and specificity (0.871) underscore the model’s strength in correctly identifying non-responders. Notably, the model’s overall AUC 0.8 confirms consistent discrimination performance at various decision thresholds. Additionally, by calculating the one-vs-all AUC for each class separately, we confirmed that the model maintains robust performance across individual classes, achieving AUCs of 0.80 for CD8 T-cell responders, 0.77 for mucosal responders, and 0.73 for systemic broad influenza A virus responders on the unseen test set (**Fig. 3C**). These values indicate that the model performs consistently well across different response types, which is crucial for a multiclass classification problem. In addition to its solid predictive metrics, the gbm model’s capacity for feature importance estimation facilitates the identification of key baseline immune predictors influencing LAIV responses. Gradient boosting methods inherently manage high-dimensional, multimodal data and effectively capture complex nonlinear relationships and feature interactions. This capability, combined with model regularization, helps prevent overfitting and ensures that the most biologically informative features retain influence in the predictive model. By systematically testing a wide array of ML approaches, we minimized biases inherent in any single algorithmic approach and ensured that our final model choice was data-driven. In summary, the gbm model’s strong predictive performance, combined with its interpretability and suitability for handling complex immunological data, makes it a powerful tool for multiclass classification of LAIV response types. Its selection underscores the importance of comprehensive model exploration and the critical role that advanced ML methods play in immunology research.

Building upon the strong predictive performance of the gbm model, we next sought to identify the baseline immune features most critical for accurately classifying individuals into CD8 T cell, mucosal, or systemic, broad influenza A virus responder phenotypes (**Fig. 3D**). The variable importance scores were derived by assessing how much each feature reduced prediction error throughout the ensemble of boosting steps, normalizing the resulting values so the top-ranked predictor received a score of 100. Features with higher scores exerted a stronger influence in distinguishing between response categories, thus offering a more nuanced and data-driven perspective than earlier studies focused on fewer or less diverse immune parameters. Our results revealed a multifaceted baseline immune landscape, extending beyond an exclusive reliance on systemic antibody levels. The top predictor was the baseline HAI geometric mean titer against H3N2 (score 100), indicating that pre-existing systemic immunity drives the type of response to LAIV observed. However, systemic factors alone did not fully explain response variability. High baseline mucosal IgA against various influenza virus antigens, including influenza B/Victoria/2/87-like lineage HA and NA (61), pH1N1 HA (48), N1 (20), H3N2 NA (18), and cH7/3 IgA (44), underscored that mucosal antibody presence at the vaccination site is also pivotal. Similarly, N1 (25) and N2 (44) titers, as well as cH6/1 ADCC (30), pointed to the importance of stalk-specific and neuraminidase-directed antibody responses, suggesting that a broad antibody repertoire can influence outcome categories. Together, these findings highlight the complementary roles of systemic and mucosal immunity in shaping the trajectory of LAIV-induced responses. Expanding the conventional view further, key cellular parameters also surfaced as prominent predictors, such as IFN-γ-producing T cells directed against influenza virus antigens (e.g., influenza A virus M/NP CD4 IFN-γ, score 64; H1N1 and H3N2 HA CD4 IFN-γ, 33 and 36; influenza B HA CD8 IFN-γ, 21) and TFH cell frequencies (34). Additionally, innate immune cells (intermediate and nonclassical monocytes, 48 and 18) and environmental factors, such as pneumococcal carriage (36) and asymptomatic respiratory virus detection (6) emerged as critical modulating factors, suggested that baseline innate immune cell landscape and subclinical microbial exposures may modulate local mucosal conditions and, in turn, influence vaccine responsiveness. Most notably, our analysis unveiled that tissue-level processes, as captured by multiple baseline nasal RNA-derived GO pathways, contribute significantly to LAIV immunogenicity. These pathways, encompassing purine metabolism (GO:0072521, 46), epithelial morphogenesis (GO:0060562, 40), Hedgehog signaling (GO:0008589, GO:0007224, scores 22 and 6), vascular and glial development (GO:0048844, GO:0021782, scores 15 and 13), fatty acid metabolism (GO:0006633, score 11), and DNA damage responses (GO:2001020, score 7), offered insights into how baseline tissue-level processes and regulatory networks may set the stage for the host’s immunological response to LAIV. The inclusion of these pathways points to a more context-dependent model of vaccine responsiveness, where tissue integrity, metabolic balance, and signaling environments can shape how humoral, cellular, and mucosal immunity come together. Collectively, these observations suggest that LAIV response phenotypes arise not from a single dominant factor but emerge from a finely tuned network of systemic and local immunity, innate and adaptive cellular components, pathogen-driven cues, and underlying tissue-level processes, ultimately guiding the host’s trajectory toward CD8 T-cell, mucosal, or systemic-broad influenza A virus responder phenotypes.

In conclusion, by employing an ensemble-based gradient boosting approach on a high-dimensional dataset, we not only demonstrated robust predictive performance in classifying distinct LAIV responder phenotypes but also uncovered previously unrecognized tissue-level and metabolic pathways that, together with systemic and mucosal immunity, shape vaccine responsiveness.

### Identifying pre-existing immune landscapes that shape LAIV responses

To delineate the pre-existing immune landscapes that define distinct immunophenotypic groups, we hypothesized that specific baseline immune conditions and mechanisms uniquely characterize each responder class. To test this, we combined ML-derived insights with comprehensive exploratory analyses of baseline seropositivity, vaccine virus shedding patterns, and detailed immunologic profiling, including baseline nasal transcriptional pathways, influenza virus-specific antibody and T-cell responses, immune cell subset frequencies, and factors such as *S. pneumoniae* load from individuals classified as CD8 T-cell responders, mucosal responders, or systemic broad influenza A virus responders. When integrated with ML-derived insights and variable importance scores, these findings underscored the biological relevance of the identified features. The resulting patterns suggest that historical exposure to influenza strains plays a pivotal role in shaping the nature of the immune response following LAIV. Subsequent analyses of seropositivity and viral shedding dynamics further support this hypothesis, providing a mechanistic understanding of how baseline immune conditions influence vaccine-induced immunogenicity (**Fig. 4**).

**Figure 4.**
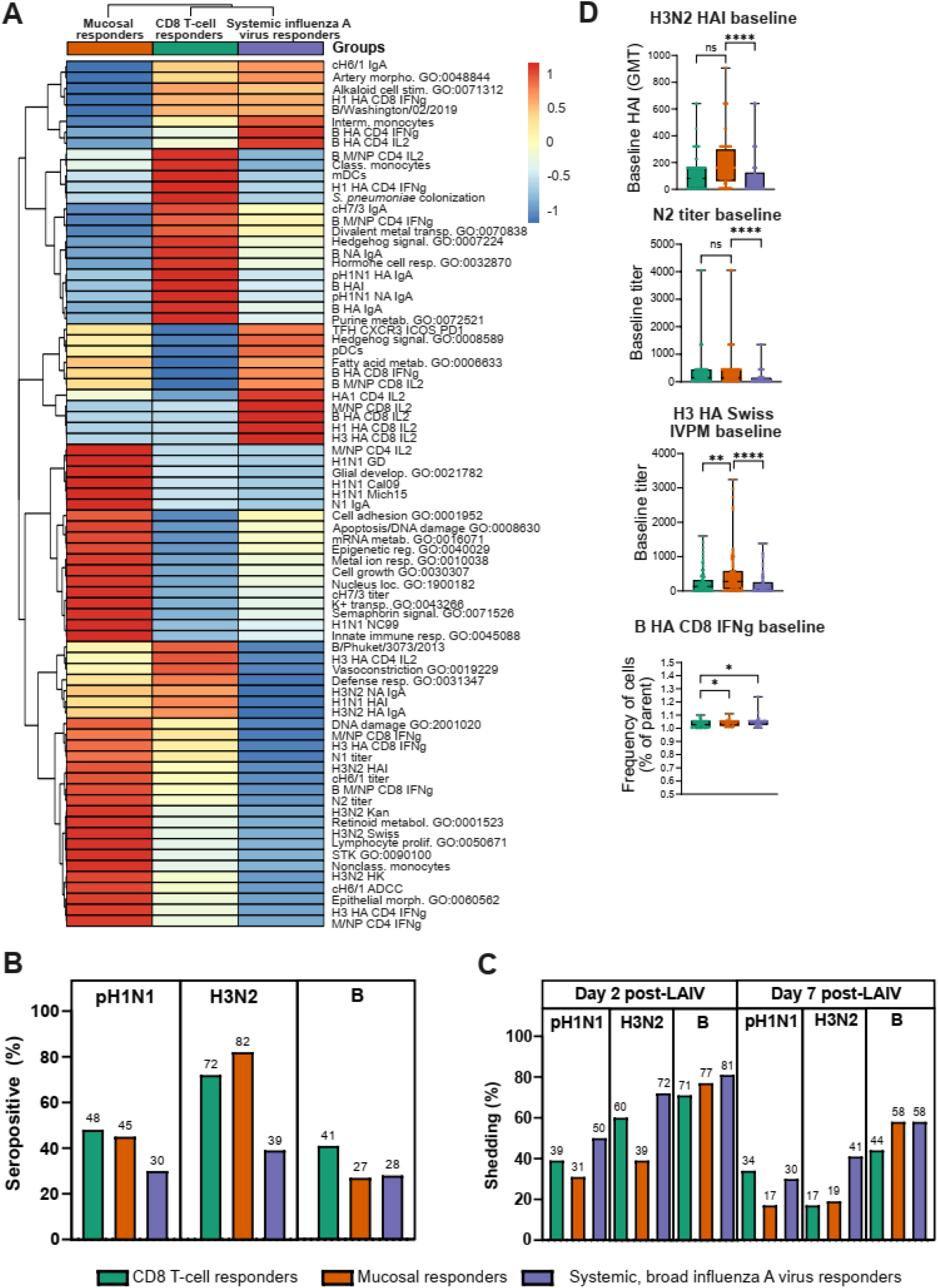
Baseline immune landscape and viral shedding profiles predictive of LAIV response groups. **(A)** Heatmap of baseline immune features predictive of LAIV response groups, organized by hierarchical clustering to show feature relationships and variations across groups (Euclidean distance, Ward’s D2 clustering method). Each cell reflects a scaled expression level, with red representing high expression and blue indicating low expression, revealing the distribution of immune features at baseline across the identified immunophenotypic clusters. **(B)** The proportion of seropositive children (HAI titer ≥10) at baseline (before vaccination) within each responder group and across all three LAIV-strains, pH1N1, H3N2, and influenza B virus (B). **(C)** The proportion of children that shed LAIV-strains (pH1N1, H3N2 and B) on day 2 and day 7 post-vaccination across all three responder groups. **(D)** Box plots showing baseline features, including H3N2 HAI geometric mean titer (gmt), titer of antibodies binding H3 HA from A/Switzerland/9715293/2013 analyzed by influenza virus protein microarray (H3 HA SWISS IVPM), titer of antibodies binding NA from group 2 (N2) and frequency of influenza B virus HA-specific CD8 T cells producing IFNγ across all three responder groups. CD8 T-cell responders (green), mucosal responders (orange) and systemic, broad influenza A virus responders (purple). Box plots denote min to max values, and points are all individuals within the group, with significance levels calculated using one-way ANOVA Kruskal-Wallis test with Dunn’s multiple comparison test to adjust for multiple testing. Significance is indicated as follows: ns = not significant, *p < 0.05, **p < 0.01, ***p < 0.001, ****p < 0.0001.

Children who develop robust CD8 T-cell responses following LAIV administration exhibit a distinctive baseline immunological and microbial signature that differentiates them from those who mount strong antibody-driven responses (**Fig. 4A**). Before vaccination, these individuals show evidence of extensive prior influenza exposures, as indicated by higher baseline seropositivity against multiple vaccine strains (as defined by HAI titers of 1:10 or greater). 41% of CD8 T-cell responders are seropositive for influenza B virus, while 48% and 72% are seropositive for pH1N1 and H3N2, respectively - proportions exceeding those observed in mucosal or systemic antibody-focused responder groups (**Fig. 4B**). Additionally, these individuals have elevated HAI responses to both H1N1 and influenza B virus, including B/Phuket/3073/2013 (B/Yamagata/16/88-like lineage), selected as a predictor in our ML model with a score of 35. This broad pre-existing seropositivity suggests a memory landscape shaped by repeated antigenic stimulation, likely due to past infections or exposures. In addition to systemic indicators of prior immunity, these children harbor enhanced mucosal defenses. They demonstrate elevated influenza virus-specific IgA levels in the nasal mucosa, targeting HA and NA from multiple LAIV strains and even chimeric group 2 (cH7/3) stalk constructs (e.g., influenza B/Victoria/2/87-like lineage NA IgA and HA IgA both scoring 61, pH1N1 NA IgA at 48, cH7/3 IgA at 44) (**Fig. 4A**), features that the ML model identified with high variable importance scores, highlighting their predictive power. IgA is a first-line defense at the respiratory epithelium, reducing viral entry and early replication. Functionally, these baseline conditions correlate with altered viral shedding dynamics after LAIV (**Fig. 4C**). While all groups initially shed influenza B virus at similar levels on day 2, by day 7 the CD8 T-cell responders show a modestly reduced shedding rate (44% versus 58% in other groups). Although pH1N1 shedding is relatively low on day 2 and changes little by day 7, H3N2 shedding declines dramatically from 60% to 17%. These patterns imply that existing mucosal immunity, particularly influenza virus-specific IgA-mediated barriers, helps limit viral replication early on. This controlled replication may provide an ideal antigenic environment for reactivating influenza virus-specific memory T-cells, allowing their rapid proliferation and functional differentiation without requiring extensive new antibody induction. Baseline transcriptional analyses further substantiate this interpretation. Key pathways, such as purine metabolism (GO:0072521; score 46) and defense response (GO:0031347; score 16), suggest a metabolic and innate immune landscape primed for rapid mobilization of memory T-cells (**Fig. 4A**). The enhanced gene signature related to the regulation of defense mechanisms underlines the ongoing interplay with microbial stimuli driving the expression of pathogen-recognition receptors, antiviral interferon-stimulated genes, antimicrobial peptides, and pro-inflammatory cytokines that collectively create a local milieu supportive of efficient T-cell recall responses. Notably, these children had elevated baseline *S. pneumoniae* loads (score 36), which can contribute to this heightened gene signature of regulation of defense response. Pneumococcal colonization has been shown to modulate mucosal immunity by inducing pro-inflammatory cytokines and enhancing antigen presentation, creating a baseline mucosal environment primed for rapid immune activation upon LAIV administration^38^. Moreover, a slightly higher proportion of children in this group had more asymptomatic respiratory viruses detected before vaccination, indicative of asymptomatic rhinovirus infection (**Fig. S2**). Rhinovirus infections induce robust innate immune responses, including type I interferon production, which enhances mucosal immunity and antiviral defense^8^. Furthermore, the Hedgehog signaling (GO:0007224, score 6) pathway was enriched at the baseline. This pathway influences T-cell differentiation and function, potentially fostering a cellular environment that supports balanced T-cell subsets and efficient recall responses^43^. Similarly, pathways linked to hormone cell response (GO:0032870), divalent metal transport (GO:0070838), and vasoconstriction (GO:0019229) may shape a tissue milieu conducive to nutrient delivery, cellular trafficking, and regulated inflammation. The elevated baseline frequency of classical monocytes and mDCs in T-cell responders adds another layer of support, as these antigen-presenting cells can efficiently capture and present influenza antigens to T-cells, thereby reinforcing the conditions necessary for prompt memory T-cell reactivation in an environment already enriched with pathogen-sensing and innate immune cues. By ensuring effective antigen presentation and co-stimulation, these myeloid cells help sustain a state of heightened immune readiness that, together with pre-existing IgA and systemic memory, promotes a swift T-cell-dominated response following LAIV challenge. Taken together, these findings suggest that children who develop CD8 T-cell-dominated responses to LAIV start from a baseline condition of extensive pre-existing influenza virus immunity, both systemically and at the mucosal surface. They have established memory B- and T-cell pools, ongoing mucosal stimulation, and a metabolic-innate readiness that set the stage for rapid T-cell recall responses. Rather than generating new antibody repertoires, these children re-engage existing immune memory, allowing CD8 T-cells, already primed by past influenza virus exposures and sustained by a supportive nasal environment, to expand and respond efficiently following the LAIV challenge.

In contrast to the CD8 T-cell responders, children who developed strong mucosal IgA responses to all three strains in the LAIV and subsequently seroconverted to influenza B virus had notably higher pre-existing immunity against influenza A viruses, including both H1N1 and H3N2 variants, but not influenza B virus (**Fig. 4A, D**). Their baseline immune landscape - especially the prominence of H3N2 HAI titers - suggests that these children had encountered influenza A viruses repeatedly, shaping a broad, cross-reactive pool of antibodies (**Fig. 4A, D**). High baseline reactivity extended beyond H3N2 to multiple H1N1 strains (e.g., A/Guangdong Maonan/SWL1536/2019, A/California/07/2009, A/Michigan/45/2015, A/New Caledonia/20/1999), neuraminidase-specific antibodies (N1, N2), and group 1 and 2 stalk-targeted constructs (cH6/1, cH7/3), collectively indicating extensive prior exposures (**Fig. 4A, D**). In addition to pre-existing influenza A virus immunity, nasal epithelial signatures pointed toward a state of readiness for rapid engagement of local immune defenses, with pathways related to epithelial morphogenesis (GO:0060562), innate immunity (GO:0045088), mRNA metabolism (GO:0016071), epigenetic regulation (GO:0040029), retinoid metabolism (GO:0001523), and lymphocyte proliferation (GO:0050671) all contributing to a stable, well-regulated mucosal interface. In practical terms, pre-existing influenza A virus immunity allowed for efficient containment of the influenza A virus strains included in the LAIV (**Fig. 4C**). These children exhibited reduced viral shedding of pH1N1 and H3N2 strains as early as day 2 post-vaccination and even more by day 7, reflecting swift local virus neutralization. However, because their baseline immunity against influenza B virus was comparatively limited (**Fig. 4B**), influenza B virus shedding remained higher and more persistent, consistent with a gap in their pre-existing influenza B virus antibody response. Still, upon exposure to LAIV, they mounted a robust humoral response against influenza B virus, ultimately seroconverting and broadening their antibody repertoire to include this lineage. In summary, these mucosal responders entered vaccination with a pre-tuned mucosal landscape and systemic antibody immunity toward influenza A virus. This configuration allowed for immediate and efficient containment of influenza A virus strains, minimizing the viral load and reducing reliance on new T-cell expansion. Their profile underscores how a background shaped by frequent influenza A virus exposure - manifested through strong baseline humoral immunity, stable epithelial architecture, and supportive innate pathways - facilitates rapid mucosal control of infection and sets the stage for subsequent seroconversion to other strains, such as influenza B virus, following LAIV.

In contrast to the CD8 T-cell and mucosal responder phenotypes, individuals who ultimately manifest a systemic, broad influenza A virus response to LAIV begin from a more immunologically naive-like baseline with respect to the vaccine strains (**Fig. 4B**). Unlike T-cell responders, who harbor extensive pre-existing immunity and IgA-mediated control, and mucosal responders, who rely heavily on robust baseline H3N2-specific immunity and a well-fortified nasal environment, these systemic responders display comparatively low baseline seropositivity across all vaccine strains (39% for H3N2, 30% for pH1N1, and 28% for influenza B virus) (**Fig. 4B**). The lack of robust preexisting humoral barriers allows for greater initial viral replication within the respiratory tract, effectively approximating a primary-like exposure that provides a more substantial antigenic stimulus (**Fig. 4C**). Shedding data confirm this scenario: systemic influenza A virus responders show higher early shedding of H1N1 (50%), H3N2 (72%), and influenza B virus (81%) on day 2 post-LAIV. As viral replication proceeds, these children gradually gain control, with shedding declining by day 7 to 30% for H1N1, 41% for H3N2, and 58% for influenza B virus (**Fig. 4C**). This improvement leads to robust seroconversion and a broadening of systemic immunity, especially against influenza A virus strains. In essence, starting from a lower baseline allows for more extensive initial viral replication, which in turn drives a strong antigenic challenge needed for de novo antibody responses. By contrast, children who become mucosal responders enter with strong pre-existing immunity focused on influenza A virus, particularly H3N2, and a mucosal environment already primed for rapid virus neutralization (**Fig. 4A, D**). As a result, they contain influenza A virus strains more efficiently, leaving them with sufficient antigenic stimulus over time to seroconvert to influenza B virus. In other words, mucosal responders use their established influenza A virus-directed immunity to quickly handle influenza A virus strains and subsequently adapt to influenza B virus, while systemic responders leverage their low baseline to mount a comprehensive, though delayed, systemic response that becomes exceptionally broad for influenza A virus but does not drive the same level of influenza B virus seroconversion. In systemic responders, baseline immunologic features, while not geared toward immediate neutralization, are nonetheless supportive of adaptive expansion. They possess a higher baseline frequency of intermediate monocytes, plasmacytoid dendritic cells (pDCs; score 15), and TFH cells (score 34) that facilitate T-cell help and B-cell maturation, fueling a robust systemic response. Additionally, systemic influenza A virus responders display a rich array of T-cell functional responses to multiple influenza HA and M/NP antigens (**Fig. 4A, D**). Although baseline antibody levels are low, the prolonged replication and higher antigen load create an environment that fosters potent T-cell priming, TFH cell activity, and the eventual generation of extensive systemic anti-influenza A virus immunity. Taken together, systemic responders exemplify how limited baseline immunity can paradoxically lead to stronger and broader systemic responses through a primary-like exposure scenario, while mucosal responders illustrate how a pre-primed environment streamlines early containment of influenza A virus, clearing the path to novel responses, including seroconversion to influenza B virus.

### The integrative machine learning interpretation approach reveals determinants of LAIV response profiles and predictors of immunogenicity

Interpreting multiclass ML models, especially those differentiating among multiple immunological outcomes, presents significant challenges due to the complexity of feature selection and the intricate interplay between features across various outcome classes. Our predictive model for LAIV response profiles required disentangling the contributions of features to three distinct responder groups: CD8 T-cell responders, mucosal responders, and systemic broad influenza A virus responders. Unlike binary classifiers, multiclass models optimize decision boundaries across multiple outcomes, making it difficult to isolate the mechanistic roles of individual features. This challenge arises because feature importance is often context-dependent, reflecting interactions, correlations, and subtle shifts in feature distributions across groups. Moreover, features selected by the ML model represent enriched and reduced characteristics across all groups, with high feature importance scores indicating either strong upregulation or significant downregulation in each group, complicating direct interpretation. To address these complexities, we adopted a multifaceted framework to extract mechanistic insights by integrating pathway-level analysis, group-specific feature impact evaluation based on model-derived contribution scores, and hierarchical examination of feature splits within the decision structure of the ML model.

To integrate and contextualize the baseline immunologic and microbiologic features influencing post-LAIV response outcomes, we conducted a pathway-level analysis as our first approach. This involved mapping each feature - from influenza-specific antibody titers and cytokine-producing T-cell subsets to nasal transcriptomic signatures and microbial loads - onto predefined biological pathways or functional categories (**Fig. 5A-D**). First, we assigned each influenza-related immune metric (e.g., H3N2 HAI titers, influenza B virus NA IgA, CD4 and CD8 T-cell cytokine responses, and IgA responses against H1N1, H3N2, and B lineages) to one of three overarching immunological compartments: ‘Humoral immunity’, ‘Cellular immunity’, or ‘Mucosal immunity’ (**Supplementary Table 2**). Separate groupings captured Microbial load (e.g., *S. pneumoniae* colonization), APC populations (e.g., pDCs, mDCs), and TFH cells due to their central role in providing B-cell help. For nasal transcriptomic data, we grouped individual GO terms into broader categories based on their biological significance. In the ‘Metabolic and Epigenetic Regulation’ category, we included pathways such as Fatty Acid Biosynthetic Process (GO:0006633), which relates to membrane synthesis and energy storage, and mRNA Metabolic Process (GO:0016071), influencing gene expression profiles. The ‘Epithelial Barrier Integrity and Tissue Remodeling’ category encompassed processes like Cell-Matrix Adhesion Pathway (GO:0001952), involved in maintaining tissue architecture, and Glial Cell Development (GO:0021782), supporting nerve and glial elements at mucosal surfaces. The ‘Immune & Inflammatory Regulation’ category included pathways such as Regulation of Defense Response (GO:0031347), fine-tuning immune effector mechanisms, and Regulation of Innate Immune Response (GO:0045088), controlling the activation threshold of innate defenses. For ‘Cell Fate & Immune Modulation’, examples included the Hedgehog Signaling Pathway (GO:0007224), guiding cellular differentiation and immune development, and ‘Epigenetic Regulation of Gene Expression’ (GO:0040029), altering the transcriptional landscape. The ‘Ion Transport/Response’ category featured processes like Metal Ion Response (GO:0010038) and Potassium Ion Transport (GO:0043266), linking ion homeostasis with epithelial barrier functions and immune signaling. Within the ‘Alkaloid Response’ category, a representative pathway was Cellular Response to Alkaloid (GO:0071312), indicating how tissues adapt to certain environmental compounds. Finally, in the ‘Stress Response’ category, the DNA Damage Response (GO:2001020) serves as an example of pathways maintaining genomic integrity and cellular resilience. When we integrated baseline features at the pathway level and computed pathway-level scores across all three LAIV-induced immunologic trajectory groups, distinct immunological signatures emerged for each phenotype. Rather than focusing on a single metric, such as a particular antibody titer or T-cell frequency, this pathway-level analysis allowed us to view the baseline immune system as a network of interacting processes. Each group’s signature emerged from the interplay of multiple biological domains, highlighting the unique baseline conditions that prime individuals for either a CD8 T-cell-dominant, mucosal antibody-focused, or systemically broad influenza A-oriented response.

**Figure 5.**
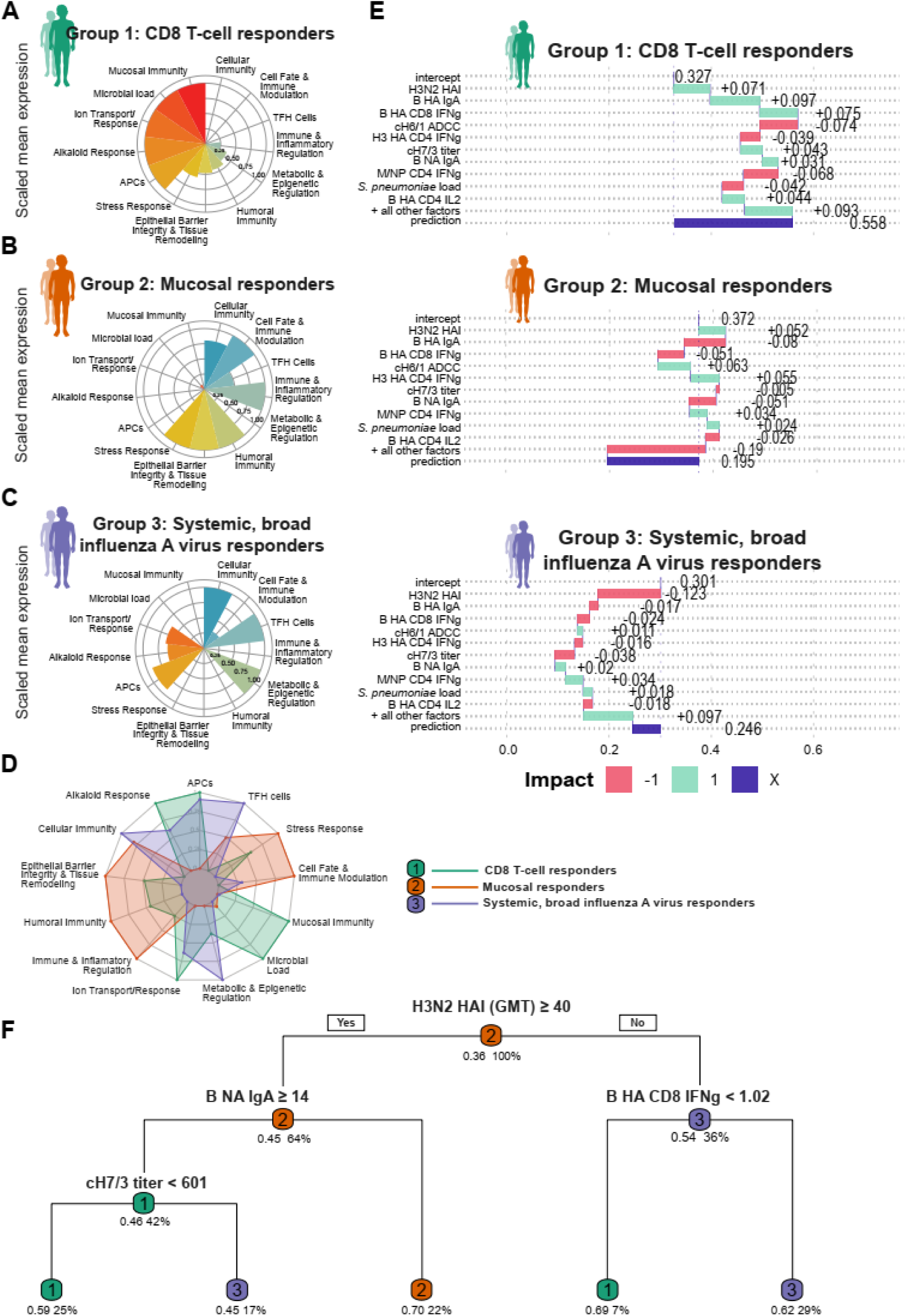
Baseline immune features and pathway-level determinants of LAIV responder profiles. **(A-C)** Polar plots illustrating scaled median expression of immune pathways across three responder groups: **(A)** CD8 T-cell responders (Group 1, green); **(B)** mucosal responders (Group 2, orange); and **(C)** systemic, broad influenza A virus responders (Group 3, purple). **(D)** Combined radar plot showing integrated immune pathway signatures across the three responder groups, highlighting inter-group differences in pathway activation. **(E)** SHAP (SHapley Additive exPlanations) summary plots showing the contribution of baseline features to model predictions for each responder group (CD8 T-cell responders, group 1, green; mucosal responders, group 2, orange; and systemic, broad influenza A virus responders, group 3, purple). The *intercept* represents the baseline prediction before feature contributions. *All other factors* include the combined effect of features not displayed in the top 10 contributors. *Prediction (purple bar)* is the final probability derived by summing the intercept, top 10 feature contributions, and all other factors. Feature impacts are color-coded: green (positive, 1) increases the likelihood of belonging to the group, and red (negative, −1) decreases it. The top 10 features are ranked by their contribution to the prediction, providing insights into key drivers of LAIV response profiles. **(F)** The decision tree depicts the splits made at each node based on immune feature thresholds. Splits are chosen to maximize class separation, with fitted class probabilities displayed as group 1 (CD8 T-cell responders, green), group 2 (mucosal responder, orange) and group 3 (systemic, broad influenza A virus responders, purple) for each terminal node. The coverage percentage represents the proportion of observations falling under each rule. Nodes are labeled with thresholds and the conditions that define group separation, with terminal nodes representing the predicted group and associated probabilities.

The dominant baseline signature for children who would later become CD8 T-cell responders included pathways related to mucosal immunity and microbial load (**Fig. 5A, D**). Other selected pathways, including those associated with ion transport/response (reflecting a mucosal environment fine-tuned by ionic conditions), alkaloid responses (potentially influenced by environmental or dietary factors), and APCs (specifically monocytes and mDCs), were also observed at baseline in children who would ultimately become systemic broad influenza A virus responders, thus are not considered a defining feature for CD8 T-cell responders (**Fig. 5D**). This baseline signature aligns with their high baseline influenza A and B virus seropositivity and robust influenza virus-specific mucosal IgA levels against all three LAIV vaccine strains, which constrain early viral replication. In addition, pre-existing microbial load due to the pneumococcal carriage continuously engages and primes the mucosal immune compartment, maintaining a baseline state of heightened innate and epithelial defense. In this environment, the presence of influenza-specific IgA at the mucosal surface acts as a front-line barrier, neutralizing the LAIV strains shortly after administration. As a result, the antigenic stimulus is insufficient to drive strong seroconversion or robust new CD8 T-cell responses, resulting instead in a modest recall of existing immunity. Together, the pre-primed influenza-specific mucosal immunity and microbial-driven baseline inflammation set a low activation threshold that efficiently contains the LAIV strains. This containment leads to less antigenic stimulation, thereby limiting both the magnitude of CD8 T-cell expansion and the impetus for producing new high-titer antibody responses. The outcome is a controlled, recall-driven response rather than a robust, de novo immunologic escalation.

The key baseline pathways in mucosal responders encompassed humoral immunity, immune and inflammatory regulation, epithelial barrier integrity and tissue remodeling, cell fate and immune modulation, and stress response (**Fig. 5B, D**). These pathways, collectively, point to a nasal environment that is well-prepared to respond adaptively, balancing protective immunity with controlled inflammation. While these individuals do not necessarily exhibit higher baseline IgA levels than other groups, they do enter vaccination with a pronounced pre-existing humoral orientation toward influenza A, particularly H3N2, as indicated by elevated H3N2 HAI titers and a broad baseline reactivity to multiple influenza A variants. In our dataset, this strong influenza A virus-focused baseline immunity correlated with reduced shedding of H3N2 and pH1N1 LAIV strains, reflecting rapid viral containment. This early control minimizes the antigenic stimulus required to mobilize *de novo* T-cell responses, allowing the immune system to reallocate resources toward countering influenza B virus. Over time, this more persistent influenza B virus replication, while initially less controlled, stimulates a robust humoral response, culminating in seroconversion to influenza B virus. The epithelial barrier-associated and immune regulation pathways identified support this concept by indicating that these mucosal responders maintain a tissue environment capable of swiftly modulating local defense and repair processes as viral replication patterns shift. Crucially, the rapid control of influenza A virus strains and the subsequent focused immune response to influenza B virus may also foster conditions for inducing a more diversified local B-cell response, thus promoting IgA class switching and affinity maturation across multiple viral antigens. In essence, the pathway-level analysis supports the conclusion that mucosal responders leverage their strong pre-existing influenza A virus immune memory to quickly manage the influenza A virus strains in the LAIV, reducing both viral load and the immediate need for extensive T-cell engagement. Therefore, they can channel their immunological machinery - guided by the tissue remodeling, immune regulatory, and metabolic cues identified by the pathway-level analysis - into mounting a successful humoral response not only against influenza B virus but also generating IgA capable of recognizing all three LAIV strains, expanding and diversifying their antibody repertoire in the process.

In children who develop systemic, broad influenza A responses, our pathway-level analysis and baseline measurements indicate that a comparatively naïve starting point - reflected in low initial seropositivity to the LAIV strains - permits more extensive early vaccine-virus replication, thereby delivering a potent antigenic stimulus. The baseline signature in this group is marked by enriched cellular immunity (multiple influenza-specific CD4 and CD8 T cells producing IFN-γ and IL-2 against all vaccine strains), APC function, and TFH cell support, conducive to vigorous germinal center reactions^44^ (**Fig. 5C, D**). Evidence from studies on inactivated influenza vaccination and acute influenza infection links day 7 activation of TFH cells expressing immune checkpoint proteins, programmed cell death protein 1 (PD-1) and inducible T-cell costimulatory (ICOS) (PD1+ ICOS+ TFH cells) with substantial increases in HAI titers^45^, underscoring the role of TFH-driven germinal center responses in shaping broad, high-affinity antibody repertoires. In this scenario, previously acquired, but not recently boosted, influenza A virus memory B and T cells can be back-boosted enhancing affinity maturation and expanding coverage to drifted and historic influenza A strains^46^. Although less pre-primed against influenza B virus, this environment strongly favors influenza A virus-centric immunity, resulting in an extensive and high-quality antibody response that transcends the boundaries of the vaccine strains.

Multiple pathways were identified in all three responder groups, but translating these broad immunologic signatures into precise, observation-level insights requires a more granular approach. For systemic responders, for instance, understanding how a relatively naïve baseline leads to a broad, drift-inclusive influenza A virus response means moving beyond generic insights about TFH- and APC-mediated germinal center reactions. We must pinpoint which specific features, such as distinct cytokine-producing T-cell subsets or particular mucosal IgA titers, exert the greatest influence under defined immunological conditions. To achieve this, we employ SHAP (SHapley Additive exPlanations) analysis^47^ (**Fig. 5E**). SHAP quantifies each predictor’s local, observation-specific contribution, revealing how each baseline variable shifts the model’s probability of classifying an individual as a CD8 T-cell responder, mucosal responder, or systemic broad influenza A responder. For CD8 T-cell responders, SHAP revealed that elevated H3N2 HAI titers and influenza B virus-directed mucosal and cellular immunity positively shifted probabilities, aligning with pathway-derived insights into pre-primed mucosal conditions and rapid recall responses (**Fig. 5E**). In mucosal responders, SHAP values highlighted the local importance of influenza B virus NA IgA, occasionally surpassing H3N2 HAI in determining the likelihood of a mucosal IgA-focused outcome, a finding consistent with their capacity to adapt pre-existing influenza A virus-biased immunity toward robust influenza B virus seroconversion (**Fig. 5E**). For systemic, broad influenza A virus responders, SHAP revealed how certain conditions, such as less pre-existing H3N2 immunity and supportive TFH-APC-metabolic landscapes, promoted the expansion of drift-inclusive antibody repertoires (**Fig. 5E**). Thus, while pathway-level analysis provided the conceptual framework, SHAP pinpointed the precise, context-dependent contributions of key immunologic factors, thereby illuminating the nuanced interplay that defines each LAIV response phenotype.

Building on the insights from pathway-level analysis and SHAP interpretation, which together identified key players and their local, context-dependent effects, we performed decision tree splits to establish explicit, data-driven cutoffs that distinguish among responder groups (**Fig. 5F**). This approach employs recursive partitioning to identify hierarchical decision rules, providing interpretable thresholds for key variables that maximize inter-group separability. By applying decision tree analysis, we identified explicit numeric cutoffs for key immunologic features that delineate each LAIV responder group. H3N2 HAI emerged as the primary discriminator (threshold of 40, **Fig. 5F**), emphasizing the central role of pre-existing H3N2-directed humoral immunity. Children exceeding this H3N2 HAI threshold and possessing moderate levels of B/Victoria/2/87-like lineage NA IgA (titer ≥14) but lower cH7/3 titers (<601) had a 59% likelihood of developing strong CD8 T-cell responses following LAIV immunization, illustrating how pre-existing H3N2 titers and constrained stalk-specific reactivity tilt the balance toward T-cell–dominated outcomes. Alternatively, a similarly high H3N2 baseline combined with low B/Victoria/2/87-like lineage NA IgA (<14) favored robust mucosal IgA and influenza B virus seroconversion (Group 2) at a 70% likelihood, indicating how variations in influenza B virus-specific mucosal immunity shift the trajectory from cellular to humoral mucosal dominance. In contrast, when baseline conditions included not only elevated H3N2 immunity and abundant influenza B virus-specific mucosal IgA but also exceedingly high cH7/3 antibody titers (≥601), the likelihood (45%) swung toward broad systemic antibody expansions against historic and drifted influenza A virus strains (Group 3). This pattern exemplifies an empirical parallel to original antigenic sin (OAS) and back-boosting: the synergistic effect of strong H3N2 reactivity and extensive group 2 stalk-targeted immunity provides the immunological substrate for broad, drift-inclusive antibody responses^46^. Importantly, our data also highlight the opposite scenario, where lower initial H3N2 HAI combined with higher baseline influenza B virus HA-specific CD8 T-cell IFN-γ responses can still produce a 62% chance of systemic breadth. Thus, while strong baseline humoral biases can drive expansive, OAS-like coverage of influenza A variants, the absence of a pronounced humoral imprint, coupled with the proper cellular conditions, can pave the way for wide-ranging antibody diversification. This underscores the flexibility of the immune system’s imprinting processes and the nuanced interplay of humoral and cellular factors shaping ultimate LAIV outcomes. Our findings provide explicit, data-driven thresholds that reveal how variations in baseline H3N2 immunity, influenza B virus-specific mucosal IgA, and group 2 stalk-targeted antibodies or cellular immunity direct LAIV-induced responses toward distinct immunophenotypes, ranging from T-cell dominance and mucosal IgA focus to broad systemic coverage, thus reflecting the complex interplay of original antigenic sin, immunodominance, and cellular support in shaping influenza vaccine outcomes.

## Discussion

In this study, we developed a novel integrative machine learning approach, *immunaut*, to unravel how preexisting immunological landscapes shape the multifaceted responses to live-attenuated influenza vaccination (LAIV). This work establishes that vaccine outcomes are not governed by singular biomarkers but emerge from a complex interplay of systemic, mucosal, cellular, and transcriptional nasal conditions. Whereas previous studies have relied on simplified linear models and focused on discrete parameters, such as HAI titers or isolated T-cell subsets, our integrative methodology allowed us to synthesize diverse data streams into a cohesive, high-resolution understanding of LAIV-induced immunophenotypes.

Our analysis extends beyond traditional paradigms by incorporating humoral, cellular, mucosal, and transcriptomic features at baseline. We identified three distinct immunophenotypic groups: CD8 T-cell responders, mucosal (and humoral influenza B virus) responders, and systemic broad influenza A virus responders and defined the precise mechanistic signatures that guide each trajectory. Coupling *immunaut’s* dimensionality reduction and clustering capabilities with pathway-level analyses, SHAP-based local feature interpretation, and decision-tree splits, we translated these high-dimensional baseline profiles into actionable immunological insights. For example, we uncovered that modest variations in H3N2 baseline immunity, influenza B virus-specific mucosal IgA, and stalk-focused antibodies serve as critical inflection points directing the immune system toward either T-cell-dominated responses, mucosal IgA-centered outcomes, or broad systemic reactivity against drifted influenza A virus strains. These findings provide data-driven support for concepts like original antigenic sin, immunodominance, and back-boosting, grounding them in quantifiable, mechanistically interpretable terms.

Crucially, *immunaut* overcomes limitations inherent in conventional analytical tools. Rather than imposing linear assumptions or isolating single parameters, it employs advanced mathematical techniques to map high-dimensional immune features into a lower-dimensional representation that preserves essential data structures. This capability reveals distinct immunophenotypic profiles without relying on prior assumptions regarding data distributions or response patterns. An illustrative example is the identification of mucosal responders who display robust mucosal IgA responses to both influenza A and B virus strains and systemic seroconversion to influenza B virus. Conventional analyses focusing strictly on systemic antibodies might mistakenly label such individuals as underprotected against influenza A virus. Yet, our integrative, nonlinear modeling clarifies that these individuals possess strong mucosal immunity to influenza A virus, likely conferring meaningful protection. This nuanced insight, reflecting complex interactions across multiple immune compartments, would be easily missed by more traditional, single-dimensional approaches.

Beyond influenza, this methodology is generalizable. *Immunaut’s* flexible design and scalability make it applicable to other vaccines and infection scenarios, enabling the incorporation of proteomic, metabolomic, and microbiome data. By providing predictive accuracy with smaller datasets and enabling the discovery of hidden patterns in immune responses, *immunaut* supports more rational vaccine design, more efficient clinical testing, and ultimately, a move toward precision vaccinology. Instead of the one-size-fits-all approach, we can now tailor vaccination strategies based on individual immune landscapes, potentially improving both efficacy and safety.

In summary, our data-driven integrative approach closes significant gaps in traditional vaccine immunogenicity studies by providing a comprehensive, mathematically robust framework for analyzing and predicting complex immune responses. This advancement is a pivotal step toward realizing the full potential of precision vaccinology - being one of the first such approaches for influenza - and paves the way for more efficient and effective vaccine development strategies.

## Acknowledgments

We sincerely thank all the participants and their families for their invaluable contribution to this study. We also thank the clinical and laboratory staff at the Medical Research Council (MRC) Unit The Gambia at the London School of Hygiene & Tropical Medicine for their dedicated support in sample collection and processing. The original study and data generation was funded by a Wellcome Trust Intermediate Clinical Fellowship award (to TIdS; 110058/Z/15/Z), the Bill and Melinda Gates Foundation (INV-004222, TIdS), an NIAID grant R21 AI151917 (FK), and the Human Infection Challenge for Vaccine development (HIC-Vac, TIdS) consortium (funded by the Wellcome Trust, UKRI MRC, and Global Challenges Research Fund (GCRF)). This research was funded by the National Institute of Allergy and Infectious Diseases (NIAID)’s Centers of Excellence for Influenza Research and Response (CEIRR) Network (AT). We acknowledge the collaborative efforts of our colleagues within the CEIRR, whose constructive feedback was instrumental in advancing this work.

## Author contributions

Conceptualization: IT, AT

Methodology: IT, SH, YJJ, BL, KH, CP, AM, JMCQ, KS, AT

Investigation: IT, SH, AT

Data curation: IT, AT, YJJ, BL, TIdS

Software: IT

Visualization: IT, AT, SH

Funding acquisition: AT

Project administration: AT

Supervision: AT

Writing – original draft: IT, SH, AT

Writing – review & editing: IT, SH, BL, YJJ, KH, AM, JMCQ, KS, FK, BK, DB, CP, AGCM, HN, TIdS, AT

## Supplementary data

**Supplementary Figure 1.**
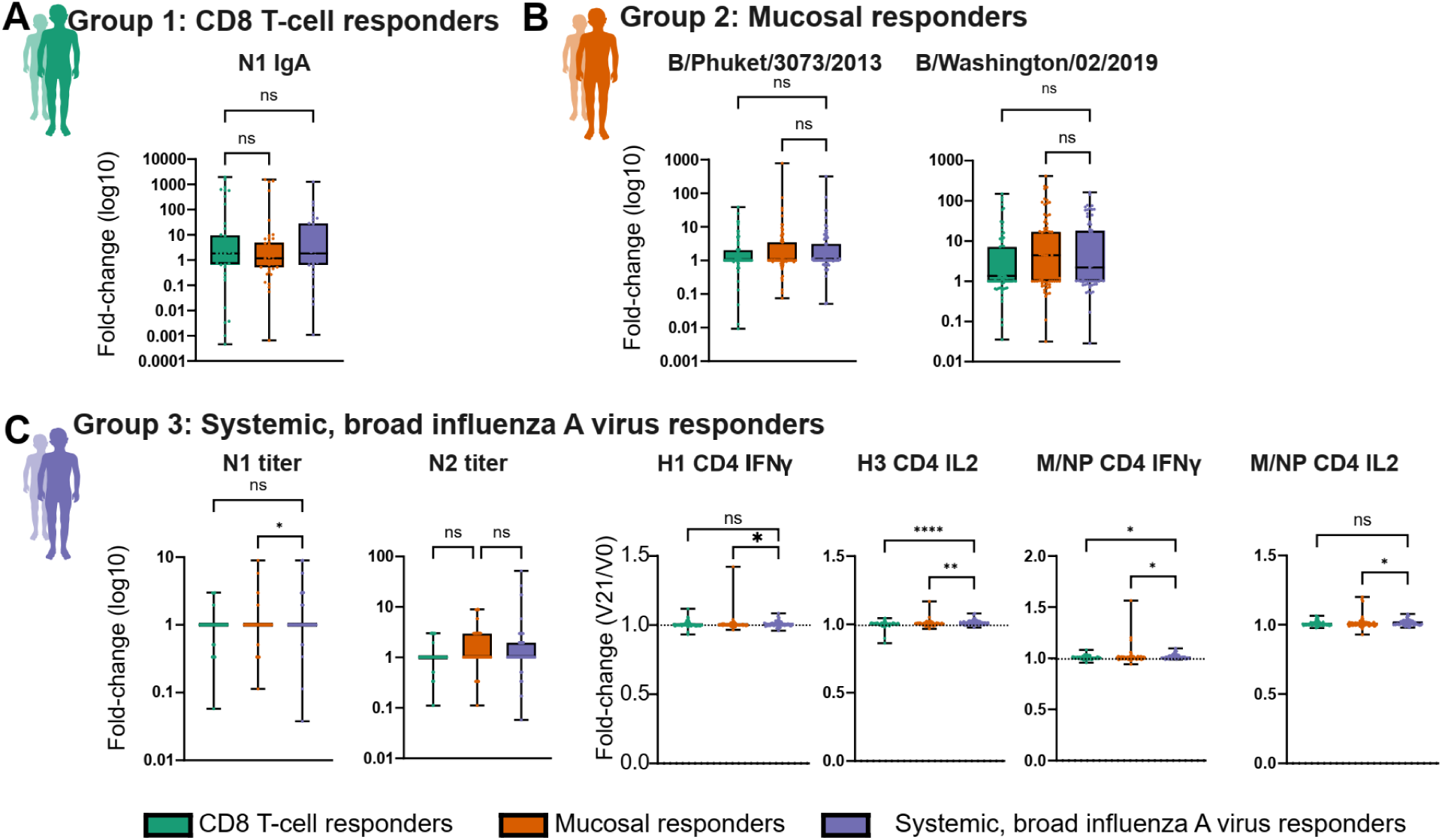
Differential immune responses across LAIV-induced responder groups. **(A)** Group 1: CD8 T-cell responders (green) distinctive feature shown as box plot fold-change in N1-specific IgA levels (log10). **(B)** Group 2: Mucosal responders (orange) show fold-changes in influenza B virus-specific IgG levels (log10) (B/Phuket/3073/2013, B/Washington/02/2019) across groups. **(C)** Group 3: Systemic, broad influenza A virus responders demonstrate fold-changes (log10) in titers of antibodies binding to N1 and N2, and fold-change CD4 T-cell cytokine responses (IFNγ and IL2) to influenza A virus hemagglutinin (H1 and H3) and matrix/nucleoprotein antigens (M/NP). Box plots denote min to max values, and points are all individuals within the group, with significance levels calculated using one-way ANOVA Kruskal-Wallis test with Dunn’s multiple comparison test to adjust for multiple testing. Significance is indicated as follows: ns = not significant, *p < 0.05, **p < 0.01, ***p < 0.001, ****p < 0.0001.

## References

1. Plotkin, S.A. & Plotkin, S.L. The development of vaccines: how the past led to the future. Nature reviews. Microbiology 9, 889–893 (2011). 10.1038/nrmicro2668

2. Hagan, T., Nakaya, H.I., Subramaniam, S. & Pulendran, B. Systems vaccinology: Enabling rational vaccine design with systems biological approaches. Vaccine 33, 5294–5301 (2015). 10.1016/j.vaccine.2015.03.072

3. Pulendran, B. & Davis, M.M. The science and medicine of human immunology. Science 369 (2020). 10.1126/science.aay4014

4. Tomic, A., Pollard, A.J. & Davis, M.M. Systems Immunology: Revealing Influenza Immunological Imprint. Viruses 13 (2021). 10.3390/v13050948

5. Sridhar, S., Brokstad, K.A. & Cox, R.J. Influenza Vaccination Strategies: Comparing Inactivated and Live Attenuated Influenza Vaccines. Vaccines (Basel) 3, 373–389 (2015). 10.3390/vaccines3020373

6. Mohn, K.G., Smith, I., Sjursen, H. & Cox, R.J. Immune responses after live attenuated influenza vaccination. Human vaccines & immunotherapeutics 14, 571–578 (2018). 10.1080/21645515.2017.1377376

7. Lindsey, B.B. et al. Effect of a Russian-backbone live-attenuated influenza vaccine with an updated pandemic H1N1 strain on shedding and immunogenicity among children in The Gambia: an open-label, observational, phase 4 study. Lancet Respir Med 7, 665–676 (2019). 10.1016/S2213-2600(19)30086-4

8. Costa-Martins, A.G. et al. Prior upregulation of interferon pathways in the nasopharynx impacts viral shedding following live attenuated influenza vaccine challenge in children. Cell Rep Med 2, 100465 (2021). 10.1016/j.xcrm.2021.100465

9. Peno, C. et al. The effect of live attenuated influenza vaccine on pneumococcal colonisation densities among children aged 24-59 months in The Gambia: a phase 4, open label, randomised, controlled trial. Lancet Microbe 2, e656–e665 (2021). 10.1016/S2666-5247(21)00179-8

10. Poland, G.A., Ovsyannikova, I.G. & Kennedy, R.B. Personalized vaccinology: A review. Vaccine 36, 5350–5357 (2018). 10.1016/j.vaccine.2017.07.062

11. Zimmermann, P. & Curtis, N. Factors That Influence the Immune Response to Vaccination. Clin Microbiol Rev 32 (2019). 10.1128/CMR.00084-18

12. Ciabattini, A. et al. Vaccination in the elderly: The challenge of immune changes with aging. Semin Immunol 40, 83–94 (2018). 10.1016/j.smim.2018.10.010

13. Plotkin, S.A. Correlates of protection induced by vaccination. Clinical and vaccine immunology : CVI 17, 1055–1065 (2010). 10.1128/CVI.00131-10

14. Pulendran, B., Li, S. & Nakaya, H.I. Systems vaccinology. Immunity 33, 516–529 (2010). 10.1016/j.immuni.2010.10.006

15. Nakaya, H.I. et al. Systems biology of vaccination for seasonal influenza in humans. Nature immunology 12, 786–795 (2011). 10.1038/ni.2067

16. Querec, T.D. et al. Systems biology approach predicts immunogenicity of the yellow fever vaccine in humans. Nature immunology 10, 116–125 (2009). 10.1038/ni.1688

17. Tomic, A. et al. SIMON, an Automated Machine Learning System, Reveals Immune Signatures of Influenza Vaccine Responses. Journal of immunology 203, 749–759 (2019). 10.4049/jimmunol.1900033

18. Tsang, J.S. et al. Global analyses of human immune variation reveal baseline predictors of postvaccination responses. Cell 157, 499–513 (2014). 10.1016/j.cell.2014.03.031

19. Krammer, F., et al. Influenza. Nat Rev Dis Primers 4, 3 (2018). 10.1038/s41572-018-0002-y

20. Koff, W.C. & Berkley, S.F. A universal coronavirus vaccine. Science 371, 759 (2021). 10.1126/science.abh0447

21. Hobson, D., Curry, R.L., Beare, A.S. & Ward-Gardner, A. The role of serum haemagglutination-inhibiting antibody in protection against challenge infection with influenza A2 and B viruses. The Journal of hygiene 70, 767–777 (1972). 10.1017/s0022172400022610

22. de Silva, T.I. et al. Comparison of mucosal lining fluid sampling methods and influenza-specific IgA detection assays for use in human studies of influenza immunity. Journal of immunological methods 449, 1–6 (2017). 10.1016/j.jim.2017.06.008

23. Meade, P., Latorre-Margalef, N., Stallknecht, D.E. & Krammer, F. Development of an influenza virus protein microarray to measure the humoral response to influenza virus infection in mallards. Emerg Microbes Infect 6, e110 (2017). 10.1038/emi.2017.98

24. Peno, C. et al. Interactions between Live Attenuated Influenza Vaccine and the Nasopharyngeal Microbiota Among Children Aged 24-59 Months in the Gambia: A Phase IV Open Label, Randomised Controlled Clinical Trial. Lancet Microbe (2023). 10.2139/ssrn.4349682

25. van der Maaten, L. & Hinton, G. Visualizing Data using t-SNE. J Mach Learn Res 9, 2579–2605 (2008)

26. Blondel, V., Guillaume, J.L. & Lambiotte, R. Fast unfolding of communities in large networks: 15 years later. J Stat Mech-Theory E 2024 (2024). ARTN 10R00110.1088/1742-5468/ad6139

27. Blondel, V.D., Guillaume, J.L., Lambiotte, R. & Lefebvre, E. Fast unfolding of communities in large networks. J Stat Mech-Theory E (2008).Artn P1000810.1088/1742-5468/2008/10/P10008

28. Tomic, A. et al. SIMON: Open-Source Knowledge Discovery Platform. Patterns (N Y) 2, 100178 (2021). 10.1016/j.patter.2020.100178

29. Biecek, P. DALEX: Explainers for Complex Predictive Models in R. J Mach Learn Res 19, 1–5 (2018)

30. Baniecki, H., Kretowicz, W., Piatyszek, P., Wisniewski, J. & Biecek, P. dalex: Responsible Machine Learning with Interactive Explainability and Fairness in Python. J Mach Learn Res 22, 1–7 (2021)

31. Tomic, A., Tomic, I. & de Silva, T. Comprehensive Multimodal Immune Response Dataset for LAIV Vaccination in Pediatric Cohorts. Zenodo (2025). 10.5281/zenodo.14719593

32. Nakajima, R. et al. Protein Microarray Analysis of the Specificity and Cross-Reactivity of Influenza Virus Hemagglutinin-Specific Antibodies. mSphere 3 (2018). 10.1128/mSphere.00592-18

33. Nachbagauer, R. et al. A chimeric hemagglutinin-based universal influenza virus vaccine approach induces broad and long-lasting immunity in a randomized, placebo-controlled phase I trial. Nature medicine 27, 106–114 (2021) 10.1038/s41591-020-1118-7

34. Kawai, A. et al. The Potential of Neuraminidase as an Antigen for Nasal Vaccines To Increase Cross-Protection against Influenza Viruses. Journal of virology 95, e0118021 (2021). 10.1128/JVI.01180-21

35. Klein, S.L. & Flanagan, K.L. Sex differences in immune responses. Nature reviews. Immunology 16, 626–638 (2016). 10.1038/nri.2016.90

36. Furman, D. et al. Systems analysis of sex differences reveals an immunosuppressive role for testosterone in the response to influenza vaccination. Proc Natl Acad Sci U S A 111, 869–874 (2014). 10.1073/pnas.1321060111

37. Fathi, A., Addo, M.M. & Dahlke, C. Sex Differences in Immunity: Implications for the Development of Novel Vaccines Against Emerging Pathogens. Frontiers in immunology 11, 601170 (2020). 10.3389/fimmu.2020.601170

38. Carniel, B.F. et al. Pneumococcal colonization impairs mucosal immune responses to live attenuated influenza vaccine. JCI Insight 6 (2021). 10.1172/jci.insight.141088

39. Mohn, K.G. et al. Live Attenuated Influenza Vaccine in Children Induces B-Cell Responses in Tonsils. The Journal of infectious diseases 214, 722–731 (2016) 10.1093/infdis/jiw230

40. Mohn, K.G. et al. Early Induction of Cross-Reactive CD8+ T-Cell Responses in Tonsils After Live-Attenuated Influenza Vaccination in Children. The Journal of infectious diseases 221, 1528–1537 (2020) 10.1093/infdis/jiz583

41. Tomic, A. et al. Divergent trajectories of antiviral memory after SARS-CoV-2 infection. Nature communications 13, 1251 (2022) 10.1038/s41467-022-28898-1

42., C.O.-M.-o.B.A.C.E.a. & Consortium, C.O.-M.-o.B.A. A blood atlas of COVID-19 defines hallmarks of disease severity and specificity. Cell 185, 916–938 e958 (2022). 10.1016/j.cell.2022.01.012

43. Crompton, T., Outram, S.V. & Hager-Theodorides, A.L. Sonic hedgehog signalling in T-cell development and activation. Nature reviews. Immunology 7, 726–735 (2007) 10.1038/nri2151

44. Crotty, S. Follicular helper CD4 T cells (TFH). Annu Rev Immunol 29, 621–663 (2011). 10.1146/annurev-immunol-031210-101400

45. Nguyen, T.H.O., Rowntree, L.C., Chua, B.Y., Thwaites, R.S. & Kedzierska, K. Defining the balance between optimal immunity and immunopathology in influenza virus infection. Nature reviews. Immunology 24, 720–735 (2024) 10.1038/s41577-024-01029-1

46. Krammer, F. The human antibody response to influenza A virus infection and vaccination. Nature reviews. Immunology 19, 383–397 (2019) 10.1038/s41577-019-0143-6

47. Ponce-Bobadilla, A.V., Schmitt, V., Maier, C.S., Mensing, S. & Stodtmann, S. Practical guide to SHAP analysis: Explaining supervised machine learning model predictions in drug development. Clin Transl Sci 17, e70056 (2024). 10.1111/cts.70056

